# Deciphering plant-beneficial fungal interactions: Unravelling metabolic diversity that underpins communication between *Laccaria bicolor* and *Trichoderma*

**DOI:** 10.1101/2024.08.26.608944

**Authors:** Prasath Balaji Sivaprakasam Padmanaban, Pia Stange, Baris Weber, Andrea Ghirardo, Karin Pritsch, Tanja Karl, J. Philipp Benz, Maaria Rosenkranz, Jörg-Peter Schnitzler

## Abstract

With over 250 known species, the genus *Trichoderma* (Ascomycota, Hypocreaceae) is found in various soils, on plant surfaces and as plant endophytes. While *Trichoderma* species are known as mycoparasites, their antagonistic behaviour can also negatively affect other beneficial fungi, such as mycorrhizal fungi. To gain insight into the metabolic signals involved in the interactions between the ectomycorrhizal fungus (ECM) *Laccaria bicolor* (Basidomycota, Hydnangiaceae), and different mycoparasitic *Trichoderma* spp. (*T. harzianum* strains WM24a1, MS8a1 and ES8g1, and *T. atrobrunneum*), we performed *in vitro* dual-confrontation experiments. We studied the volatile organic compounds (VOCs), hyphal metabolomes and soluble metabolites released by each of the fungi in various co-cultivation scenarios. The results revealed an altered growth of the mycelia depending on the degree of contact: When *Trichoderma* spp. and *L. bicolor* shared only the same headspace, *Trichoderma* spp. growth was at least partially inhibited, whereas in direct contact the growth of *L. bicolor* was impaired. Distinct strain- and species-specific changes in hyphal metabolites, in exudates and volatile emission were revealed from each of the studied fungi. We identified both core metabolite profiles and interaction-specific metabolic responses that were related to carbohydrate, lipid, nucleotide, energy and amino acid metabolisms. Volatile and soluble metabolites revealed temporal and spatial adjustments in dual cultures compared to solitary cultures, suggesting rapid contact-dependent adaptations and demonstrating the dynamic communication mechanisms between *Trichoderma* spp. and the ECM. These results suggest a central role for both emitted and secreted fungal metabolites in the fungal non-self-recognition and in interaction with each other.

## Introduction

The rhizosphere is home to a diverse range of microbial species, including bacteria, yeasts, protozoa, archaea, and fungi, all of which play an important part in soil health and plant well-being (Adeleke & Babalola, 2021; Liu et al., 2021). When microbial bioinoculants are used as biocontrol agents, competitiveness and proper establishment in the rhizosphere are prerequisites for effective biocontrol regardless of the mode of action of the biocontrol agents (BCA) used (Liu et al., 2022; Manzar et al., 2022; Raaijmakers et al., 1995; Rush et al., 2021; Weller, 1988). Among the available BCAs, fungi play multiple roles with high plasticity and adaptability to adverse and unfavourable conditions (Sun et al., 2005). Currently, *Trichoderma* spp. dominate the bioinoculant market with many of the 250 identified species showing biocontrol activities (Bissett et al., 2015; Guo et al., 2019). *Trichoderma* spp. occur in diverse environments (Olowe et al., 2022) and in complex interactions with other soil microflora (Sun et al., 2005). They can control and antagonise plant pathogenic fungi through several direct and indirect mechanisms (Asad, 2022), including competition for space and nutrients (Harman et al., 2004; Shoresh et al., 2010), secretion of various hydrolytic enzymes such as chitinases, glucanases and proteases, or emission of volatile organic compounds (VOCs) (Guo et al., 2019; Stange et al., 2024; Schulz-Bohm et al., 2017). Some effects of fungal antibiotics and effector molecules have also been reported in fungal confrontations (Chernin et al., 2011; Hartmann & Schikora, 2012; Walker et al., 2003). *Trichoderma* spp. is especially interesting BCA, as it does not only antagonize pathogens, but it also may have the potential to discriminate between plant-beneficial and pathogenic fungi (Hassani et al., 2018; Stange et al., 2024). Recently, Stange et al. (2024) showed that the growth of *Trichoderma* spp. display positive tropism towards plant pathogens and a negative tropism in the presence of ectomycorrhizal (ECM) fungi, such as *Laccaria bicolor*. In this interaction, communication could occur either via VOCs, which have been previously shown to play a role in fungal-plant and fungal-fungal interactions (Ditengou et al., 2015; Guo et al., 2019; Guo et al., 2021) or via soluble metabolites. These compounds could be important players in the mutual perception of fungi.

The term VOCs encompasses a wide range of chemically diverse small molecules released by plants, microbes and fungi (Baldwin et al., 2006; Guo et al., 2019; Šimpraga et al., 2016). To date, approximately 480 different VOCs have been detected from *Trichoderma* spp. (Dourou & La Porta, 2023; Siddiquee, 2014), whereas only 15 VOCs, mostly terpenes, have been reported from *L. bicolor* (Müller et al., 2013; Ditengou et al., 2015; Guo et al., 2019). *Trichoderma* VOCs include heterocycles, aldehydes, ketones, alcohols, phenols, thioalcohols, thioesters and their derivatives, and hydrocarbons such as monoterpenes (MT) and SQTs and their derivatives, i.e., oxygenated monoterpenes (oMT), and oxygenated sesquiterpenes (oSQT) (Dourou & La Porta, 2023; Siddiquee, 2014). Previous studies on *Trichoderma* spp. VOC production have shown that their emission profiles vary depending on the species or strains and on the cultivation environment (Stoppacher et al., 2010; Crutcher et al., 2013). The VOC profiles depended, moreover, on the contact degree with other fungi (*L. bicolor*), i.e. the emission profiles were altered depending on if the communication occurred only through headspace (aerial contact-AC), shared growth media (media contact-MC) or direct hyphal contact (direct contact-DC) (Guo et al., 2019). However, the basis of interactions between *Trichoderma* spp. added to soils and different fungi (e.g. ECM) present in the rhizosphere remains unclear.

In addition to VOCs, the confrontation between different fungal hyphae leads to changes in mycelial growth and secretion of soluble secondary metabolites (Boddy, 2000; Heilmann-Clausen & Boddy, 2005; Hu et al., 2011). Such bioactive allelochemicals are “non-nutritional chemicals produced by individuals of one species that affect the growth, health, behaviour, or population biology of another species” (Holighaus & Rohlfs, 2016). The genomes of *Trichoderma* spp. and *L. bicolor* are rich in genes encoding enzymes responsible for the production of secondary metabolites (Kubicek et al., 2011; Plett et al., 2015; Nosenko et al., 2023). These metabolites, which include, among others, signalling molecules, growth inhibitors, toxins and their by-products (Azzollini et al., 2018; Glauser et al., 2009; Heilmann-Clausen & Boddy, 2005), can contribute to a potential competitive advantage for *Trichoderma*’s biocontrol activity. Soluble metabolites may also mediate various fungal-plant interactions, for example in establishing a symbiotic relationship between *L. bicolor* and the host (Kubicek et al., 2011; Mukherjee et al., 2012). Understanding the recognition and interaction between *L. bicolor* and *Trichoderma* spp. is crucial given their co-occurrence in shared niches and rhizospheres, such as that of poplar (Guo et al., 2019; Kaling et al., 2018; Stange et al., 2024; Sivaprakasam Padmanaban et al., 2022) and the common use of *Trichoderma* spp. as BCA. As for BCA, the extent of the interaction effects depends on the duration of inoculation and the concentration of secreted allelochemicals (Zin & Badaluddin, 2020). Only in the recent years, there has been an increased focus on deciphering the chemical composition of allelochemicals released by *Trichoderma* and their effects on biochemical and physiological processes, with potential field applications (Dutta et al, 2023). Further studies are essential to understand the function of individual metabolites in the interaction of *Trichoderma* spp. with other, potentially beneficial fungi sharing the same habitat.

Therefore, in the present study, we confronted the ectomycorrhizal fungus *L. bicolor* with either of four mycoparasitic *Trichoderma* spp. - *T. harzianum* strains WM24a1, MS8a1 and ES8g1 and *T. atrobrunneum* (Guo et al., 2019,2020; Stange et al., 2024). We aimed to track changes in the dynamics of volatile and soluble metabolites to elucidate the interaction between *Trichoderma* spp. and *L. bicolor* over distance and time. We hypothesized that (I) the VOC and exudate patterns of the different fungi are strongly dependent on the degree of interaction with the other fungi leading to growth inhibition and (II) are conserved across species. Furthermore, we postulated that (III) the interacting fungi could sense each other through signalling compounds (VOCs, exudates) and subsequently alter their metabolic pathways. Finally, we hypothesized that (IV) the interacting fungi could release soluble metabolites into the interaction zone that could help defend/avoid the other fungi.

## Materials and Methods

### Fungal strains, media and cultivation

Three strains of *Trichoderma harzianum* (WM24a1, MS8a1 and ES8g1), *Trichoderma atrobrunneum* and *Laccaria bicolor* (strain S238N) were used in the study. The fungi were inoculated as mycelial plugs (1 cm diameter) in glass Petri dishes of 10 cm diameter containing 40 ml of Modified Melin-Norkrans (MMN) synthetic medium covered with a sterile cellophane sheet (as described in Müller et al., 2013). The inoculated plates were sealed with parafilm and incubated in permanent darkness at 23°C in a growth chamber.

### Experimental setup and growth analysis of fungi

*L. bicolor,* as a slow-growing species, was initially inoculated on one side of the Petri dishes for co-culture. Split Petri dishes were used to study the interactions in AC and unsplit Petri dishes for other interactions (MC and DC). After 14 days of cultivation, an agar plug of *Trichoderma* strain was inoculated on the other side of the same Petri dish. Five replicates of each *Trichoderma* strain in each interaction scenario and pure cultures (PC) were prepared and examined according to Guo et al. (2019). Images were taken at 3,5 and 7 days after co-culture using a Nikon D300 camera (60 mm Nikkor AF-S Micro-Nikkor Lens, Nikon, Tokyo, Japan) and used for growth analysis. Growth inhibition was calculated using the following formula (Raut et al., 2014):

Growth Inhibition (%) = D1−D2/D1×100

D1: fungal area grown alone.

D2: fungal area grown in co-cultivations.

Three days post *Trichoderma* inoculation, the headspace VOCs were collected from the five replicates of PC, AC and MC co-cultivation scenarios. Similarly, mycelium and media from PC, MC and DC were used for non-targeted metabolomic analysis.

### VOC profiling

VOCs were extracted from the headspace of PC, AC and DC fungal cultures for 16 h using Twisters (Gerstel GmbH & Co.KG, Mülheim an der Ruhr, Germany) and the Headspace Sorptive Extraction (HSSE) technique according to the method described by Müller et al. (2013). Subsequently, the Twisters were stored in 4°C and later subjected to VOC analysis through Thermal Desorption-Gas Chromatography-Mass Spectrometry (TD-GC-MS), adhering to protocols as previously detailed (Ghirardo et al., 2012; Ghirardo et al., 2016; Guo et al., 2021).

Putative Annotation of the mass spectra was performed by comparison with best hits from the libraries of reference spectra (NIST 11, Wiley 275). For quantification, response factors were calculated using standards: sabinene and α-pinene for MT, linalool for oMT, β-caryophyllene and α-humulene for SQT, and geraniol for oSQT. The resulting data were input for subsequent statistical analysis.

### Non-targeted metabolomic analysis

The area of mycelium and media (beneath the mycelium and cellophane) of MC, DC and PC was divided into three zones (Figs. 4 and 6). The mycelium was scraped from the cellophane with a spatula, the cellophane was removed after sampling and the media was sampled from the same area where the mycelium was collected. The collected samples were stored at -80°C for further processing and extraction. The media were freeze-dried at −50°C under 0.040 mbar vacuum (Alpha 1-4 LDplus, Christ, Osterrode, Germany).

For the extraction process, 800 µL of a cold (5°C) mixture of methanol: 2-propanol: water (in a ratio 1:1:1, v/v/v) extraction solvent was added to 25 mg of mycelium and freeze-dried media. This solvent contained 50 µL of an internal standard (IS) mixture (Supplementary S2). The samples were gently mixed in a 2 mL polypropylene tube for 1 minute. These mixtures were then ultrasonicated for 10 min in a water bath held at approximately 5°C. After sonication, the solution was centrifuged at 10*g* for 10 minutes at 4°C. This step resulted in the collection of 640 µL of supernatant. The collected supernatant was then subjected to drying using a SpeedVac system (Univapo 150H, Uniequip, Planegg, Germany). The resulting residue was reconstituted in 400 µL of a solution of 50% (v/v) acetonitrile in water. After a mixing step of 1 min and subsequent centrifugation at 10 *g* for 10 min at 5°C, 300 µL of supernatant was carefully transferred into 350 µL amber glass vials (following Bertić et al. (2021).

Untargeted metabolomics analysis was carried out following the methodologies outlined by Ghirardo et al. (2020) and Hemmler et al. (2018), Each sample underwent separate analysis using the RPLC-positive (+) and HILIC-negative (-) columns electrospray ionization (ESI) modes.

### Metabolic profiling

The LC-MS data were analysed using Metaboscape 4.0 (Bruker). This software facilitated post-acquisition processes, including peak picking, alignment, isotope filtering, and peak grouping based on peak-area correlations (as per Domingo-Almenara et al. (2018)). Detailed parameter settings are provided in Supplementary S3. Results from RP and HILIC analyses were merged manually.

Annotations were based on available MS/MS spectra matched to different libraries, including HMDB (http://www.hmdb.ca/) (Wishart et al., 2009), MoNa, Vaniya/Fiehn Natural Products Library and LC-MS/MS spectra. For the other compounds the smart formulas were used for putative annotation.

The data were normalised using the peak intensities of internal standard (IS) mixture (S2) and missing values were replaced with the random values below the detection limit (1-800). The resulting data was used for the subsequent statistical analysis.

### Statistical analysis

The growth area of the fungal mycelium was measured using the image analysis software IMAGEJ (Schneider et al., 2012). The significance of growth inhibition was tested by a one-way ANOVA followed by Tukey HSD post hoc test (p<0.05). All data were always logarithmically (log10) transformed, centred, and Pareto scaled (Bertić et al., 2021). Principal component analysis (PCA) and orthogonal partial least square regression discriminant analysis (OPLS-DA) of both VOC and metabolic data were performed in SIMCA-P v.13.0.3.0 (Umetrics, Umea, Sweden). Features with a Variable Importance of Projection (VIP) score greater than 1 were selected for the calculation of log2fold change (>1), p-value and adjusted p-value (<0.05) to identify potential discriminative mass features. Detailed processing parameters can be found at Bertić et al., 2021. Hierarchical clustering analysis (HCA) of VOC concentration and metabolic mass features with VIP>1 was used to visualise expression patterns. Enrichment- and pathway analysis of the putatively significant annotated features was performed using Metaboanalyst 5.0 (https://www.metaboanalyst.ca/MetaboAnalyst/home.xhtml) (Pang et al., 2021). The resulting pathways were cross-checked using the pathway analysis function in Metaboanalyst 5.0 (KEGG). The occurrence of the pathway function in corresponding fungi and the biological process of the pathway were manually checked on the KEGG database (https://www.genome.jp/kegg/pathway.html) (Kanehisa et al., 2023). Alluvial plots were generated using RAWGraphs 2.0 (Mauri et al., 2017). Chemical class classification of the significant compounds was performed using the ’Multidimensional Stoichiometric Compound Classification’ (MSCC) approach, based on elemental ratio compositions (Rivas-Ubach et al., 2018).

## Results

### Co-culture of *L. bicolor* and *Trichoderma* mutually limit mycelial growth and lead to altered VOC emission profiles

When incubating *L. bicolor* with *Trichoderma* spp. in co-culture (Fig. 1A), different growth inhibition dynamics could be observed (Figs. 1C, S1C, S2C and S3C). On day three, MS8a1 did not inhibit *L. bicolor* growth in AC conditions and only modest inhibition was observed in MC conditions (7.8 ± 2.8). MS8a1 growth was inhibited by 15 ± 10% in AC and 39 ± 10% in MC. By the fifth day, both species were similarly affected. MS8a1’s growth was reduced by 10 ± 7% in AC and 27 ± 9% in MC, while the growth of *L. bicolor* was reduced by 10 ± 4% in AC and 22 ± 10% in MC. By day seven, MS8a1 significantly inhibited the growth of *L. bicolor* (36 ± 10% in AC and 32 ± 8% in MC), while the ECM had little effect on MS8a1 (5 ± 2% in AC and 15 ± 8% in MC). A similar behaviour was observed with the other *Trichoderma* strains (Fig. S1C, S2C, and S3C). On day three, an AC contact with *L. bicolor* inhibited the growth of WM24a1 by 37 ± 5%, ES8g1 by 29 ± 7% and *T. atrobrunneum* by 24 ± 7%, while *L. bicolor* growth was not inhibited compared to PC. In the later stages of co-culture, *L. bicolor* growth was also inhibited by 11 - 38% (Figs. S1C, S2C and S3C). The reciprocal growth inhibition was evident in the AC scenario and was also observed in the MC conditions, albeit to a slightly greater extent.

**Fig 1.**
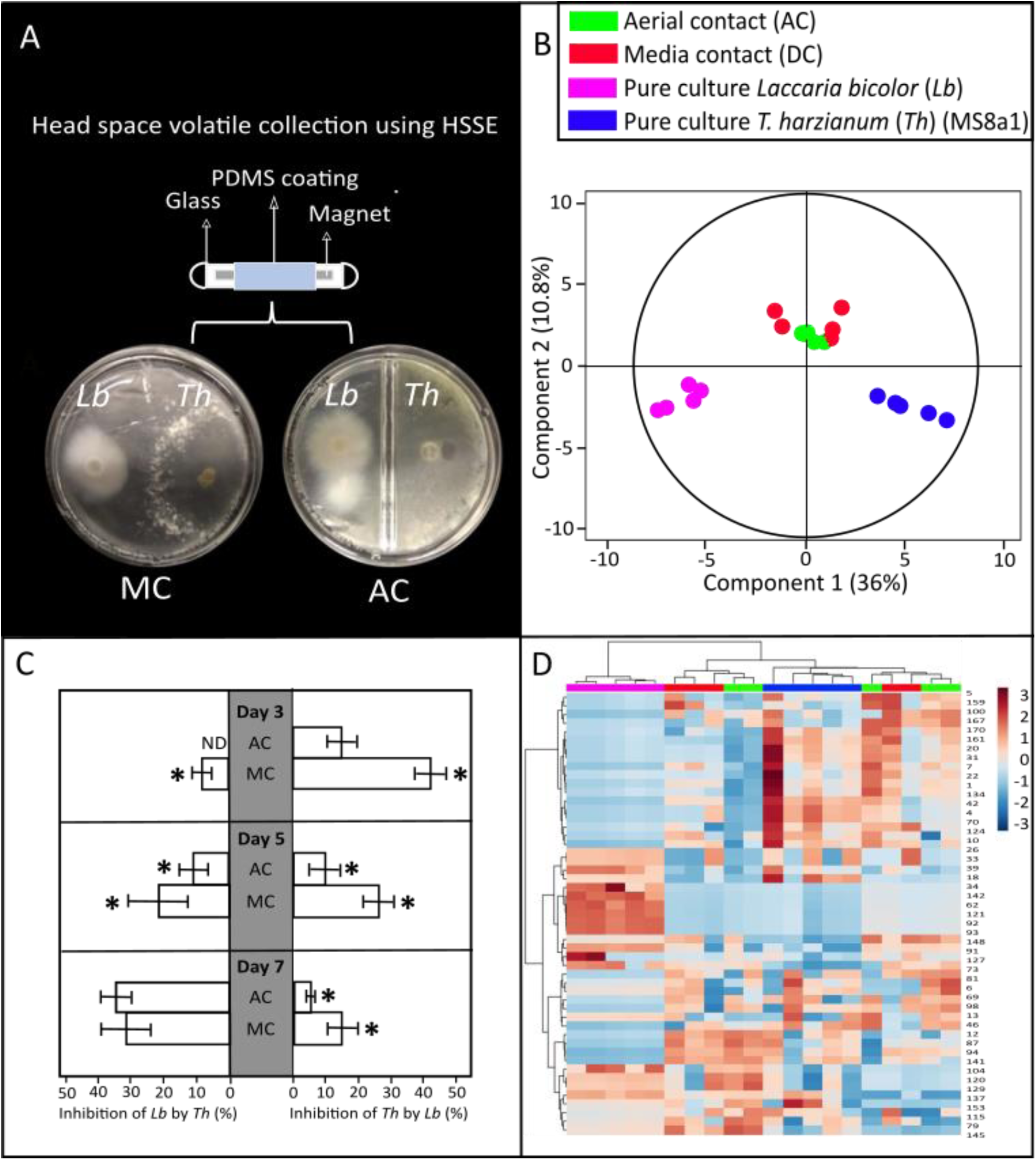
VOC analysis of *L. bicolor* (*Lb*) and *T. harzianum (*MS8a1) (*Th*) co-cultivated either in aerial contact (AC) or media contact (MC). (A) Experimental setup of the VOC collection from the fungal co-cultivation in split and non-split Petri plates. (B) Orthogonal partial least square regression discriminant analysis (OPLS-DA) showing differences among VOC profiles under different levels of co-cultivation. OPLS model fitness: R^2^X(cum) = 0.855, R^2^Y(cum) = 0.787, Q^2^Y(cum) = 0.625, CV-ANOVA=7.2 x 10^-7^. (C) Growth inhibition of *Th* on *Lb* (left) and *Lb* on *Th* (right) under different levels of co-cultivation compared to growth on pure cultures. Significances within each day are denoted as asterisks (one-way ANOVA and Tukey HSD, p < 0.05); mean ± SE; values are average of 5 replicates. (D) Two-dimensional hierarchical clustering analysis of the VOC emissions from the two fungi grown as MC, AC or pure culture. ND: not detected.

**Table. 1.**
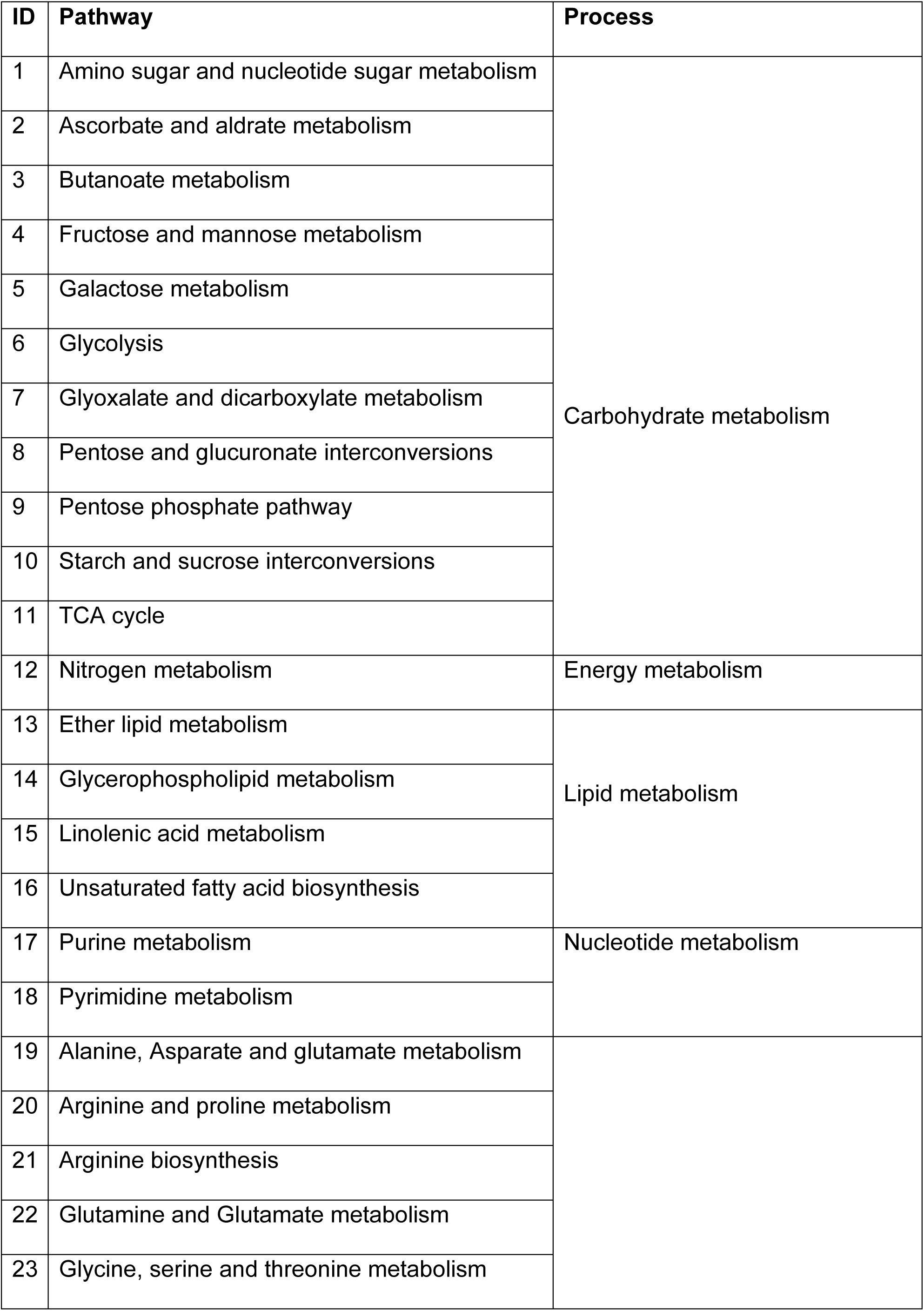
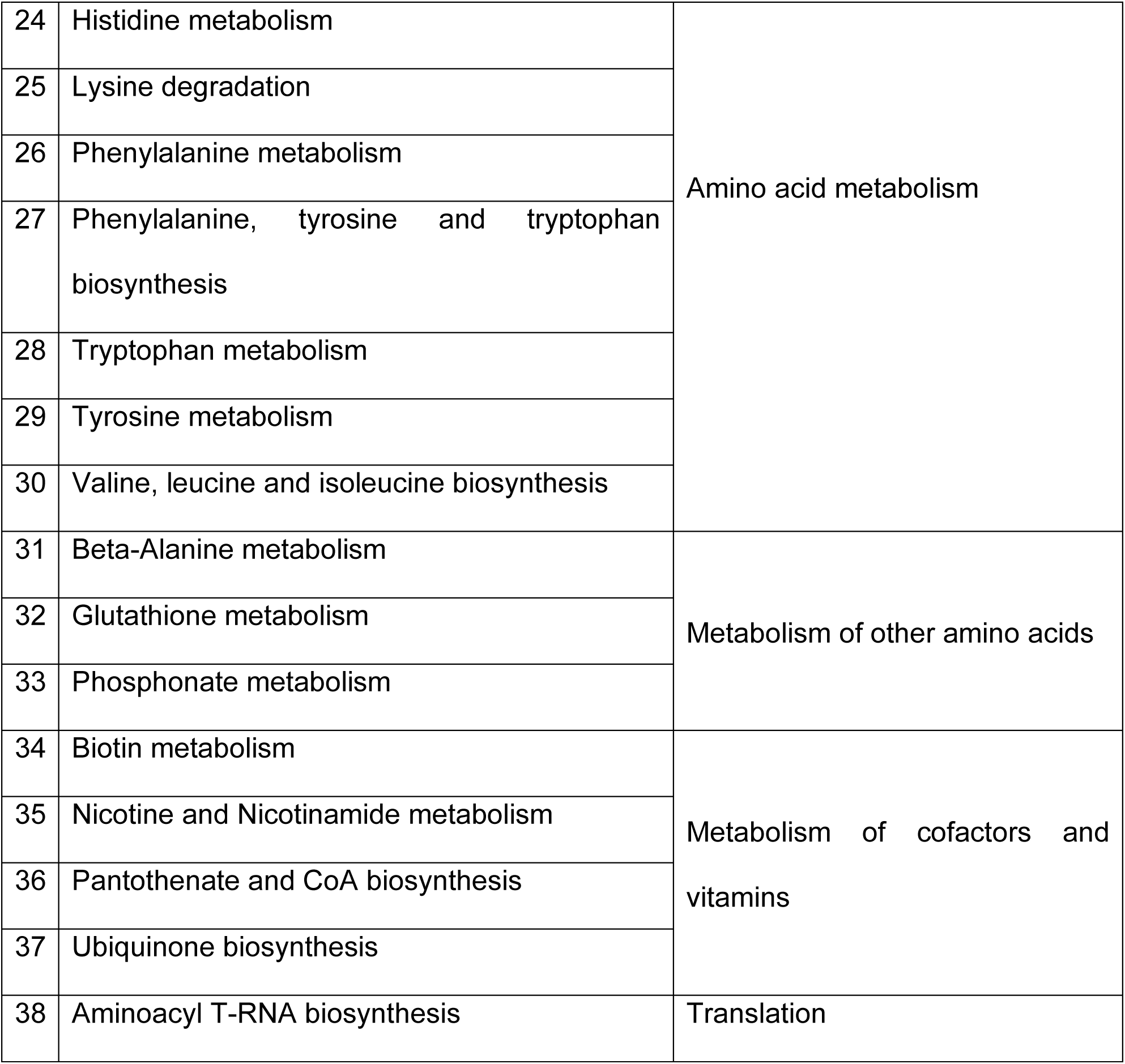
Pathways enriched by putatively annotated and significant metabolic features grouped according to metabolic processes.

The VOC analysis of *L. bicolor* and MS8a1 showed distinct profiles and emission strengths for the two species in solitary cultures, as shown by orthogonal partial least square regression discriminant analysis (OPLS-DA; CV-ANOVA = < 0.001; 36% of VOC variance (Y axis) explained; Fig. 1B). For the other *Trichoderma* strains, WM24a1, ES8g1 and *T. atrobrunneum,* the differences in VOCs detected in PC explained 21.5%, 11.5% and 7.78% of the variance, respectively with *L. bicolor* (Fig. S1B, S2B and S3B). Hierarchical clustering analysis (HCA) of VOCs from *L. bicolor* and MS8a1 in different degrees of contact revealed distinct clusters in PC compared to AC and MC (Fig. 1D). These clusters show individual, interaction-degree-specific VOC emission patterns (Figs. S1D, S2D, and S3D). Co-culture resulted in the loss of VOCs originally detected in *L. bicolor*, putatively annotated as p-cymene, γ-curcumene, nonanal, and furfuryl alcohol, and the detection of new VOCs. (Fig. 1D and Figs. S1D, S2D, S3D, Table S1).

Figure 2 (and Table S1) shows VOC emission (volatile terpene) levels and patterns for each fungus in different culture conditions: in PC, 22 compounds for WM24a1, 21 for MS8a1, 26 for ES8g1, 32 for *T. atrobrunneum*, and three for *L. bicolor.* MS8a1 showed the lowest terpene emission rate (38 pmol cm² h⁻¹), while *T. atrobrunneum* had the highest (181 pmol cm² h⁻¹). *Trichoderma* spp. emitted mainly sesquiterpenes, while *L. bicolor* emitted mainly monoterpenes. Co-culture significantly influenced VOC patterns and intensities compared to PC. AC of *L. bicolor* and *T. harzianum* (WM24a1) increased emissions, while other *L. bicolor* - *Trichoderma* combinations reduced total emissions. In general, the occurrence and loss of VOCs were highly species-specific (Fig. 2A). The core VOC profile of all fungi under all co-cultivation scenarios were 41 in WM24a1, 27 in ES8g1, 39 in MS8a1 and 50 in *T. atrobrunneum*. There were only a few lost compounds (Fig. 2B), but formation of new VOCs, i.e., those that were not found in pure cultures, were detected during co-cultivation scenarios (Fig. 2B) (see Table S1), suggesting a potential role in intraspecific communication.

**Fig 2.**
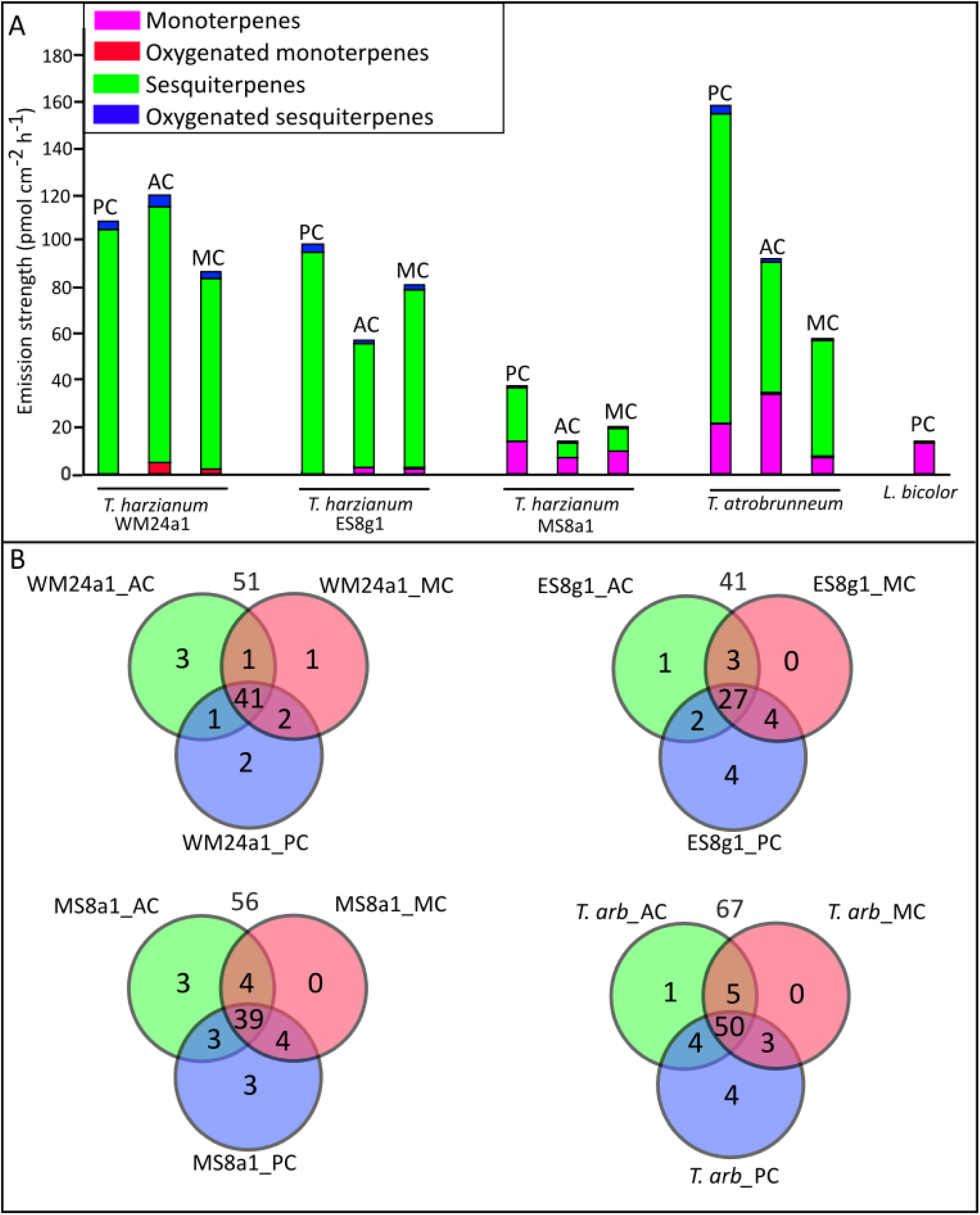
VOC profile of different fungal species. (A) The total emission of various VOCs across three *T. harzianum* strains (WM24a1, MS8a1 and ES8g1), *T. atrobrunneum (T. arb)* and *L. bicolor* grown as pure cultures (PC) and as co-cultures in aerial (AC) and media contact (MC). The VOCs are classified into four different chemical groups as monoterpenes, oxygenated monoterpenes, sesquiterpenes and oxygenated sesquiterpenes. Values are an average of 5 replicates. (B) Venn plots depicting the number of compounds detected from the different *Trichoderma* strains across different growth and interaction conditions with *L. bicolor*. The number on top of each Venn plot indicates the total number of VOCs identified.

Co-culture of *L. bicolor* and WM24a1 increased emissions and led to the generation of four new compounds during AC: propylcyclopentane, 1-cyclopentylethanone, 2-methyl-5-propylthiophene, and 4-*T*-butylphenol. In MC, α-cuprene and 4-*T*-butylphenol were newly observed. In the *L. bicolor* - ES8g1 combination, emissions were lower, but also in this case new compounds were identified: 1-(dodecyloxy)-2-nitrobenzene, methylcycloheptane, and 1-methyl-4-trimethyl-cyclobenzene in MC and β-Curcumene only in AC. The combination *L. bicolor* - MS8a1 had the lowest total emissions but led to the detection of three new VOCs in AC: heptamethyl-2-nonene, di-n-decyl ether, and 1,1,3,5-tetramethyl-cyclohexane. In the *Laccaria* - *T. atrobrunneum* co-culture, five new VOCs, including 1-methyl-4-cyclohexene, were identified exclusively in AC.

### Non-targeted metabolomics reveals strain- and species-specific metabolic diversity among *Trichoderma* and *Laccaria*

We analyzed total metabolite profiles of the hyphae and underlying media (exudates) of the pure cultures. The aim was to create reference metabolomes that could be used for comparison with the hyphal metabolomes during co-culture, to detect metabolic shifts (Fig. 3A) and the secretion of potential signalling compounds into the medium (Fig. 3B). We detected a total of 9365 mass features, of which 540 could be putatively identified (Level 2 - matched to the spectral library), 7314 tentatively annotated compounds (Level 3 - matched to the library using R codes and based on MS1 data (Bertić et al., 2021)), 1450 metabolites matched to broad compound classes based on molecular formula predicted by MS1 data (Level 4 using the Multidimensional Stoichiometric Compound Classification (MSCC) approach), and 61 mass features of so far unknown unique features (Level 5) (Bertić et al., 2021). The metabolome analysis (Fig. 3A only shows Level 1 compounds) revealed a variety of metabolites comprising different chemical classes, including amino sugars, carbohydrates, lipids, peptides and amino acids, phytochemicals, and unclassified compounds. These metabolites were widely distributed among the different fungal species. While the carbohydrate composition of the fungal hyphae showed a striking similarity, strain-dependent variations were observed for lipids, peptides and phytochemicals (Fig. 3A).

**Fig 3.**
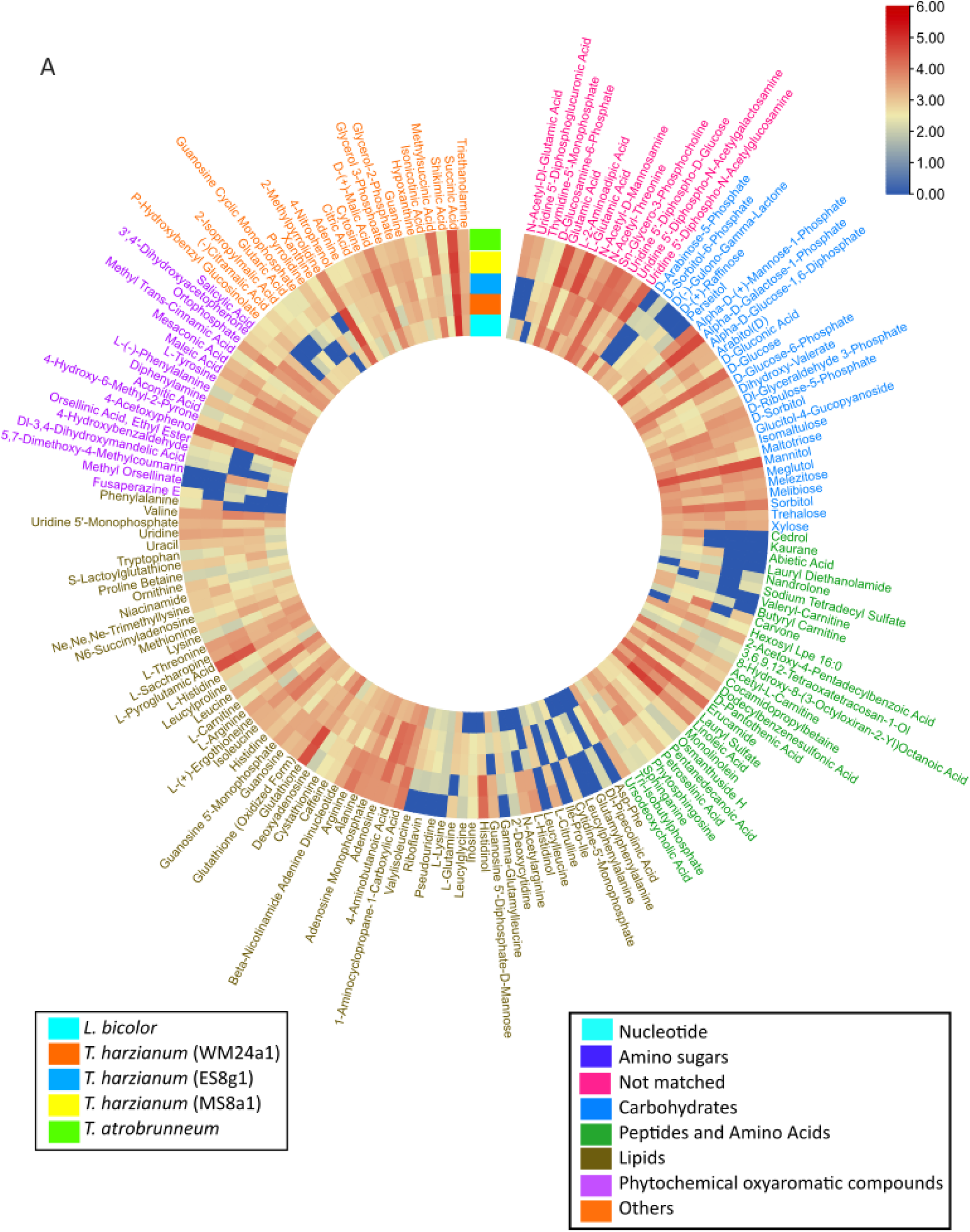
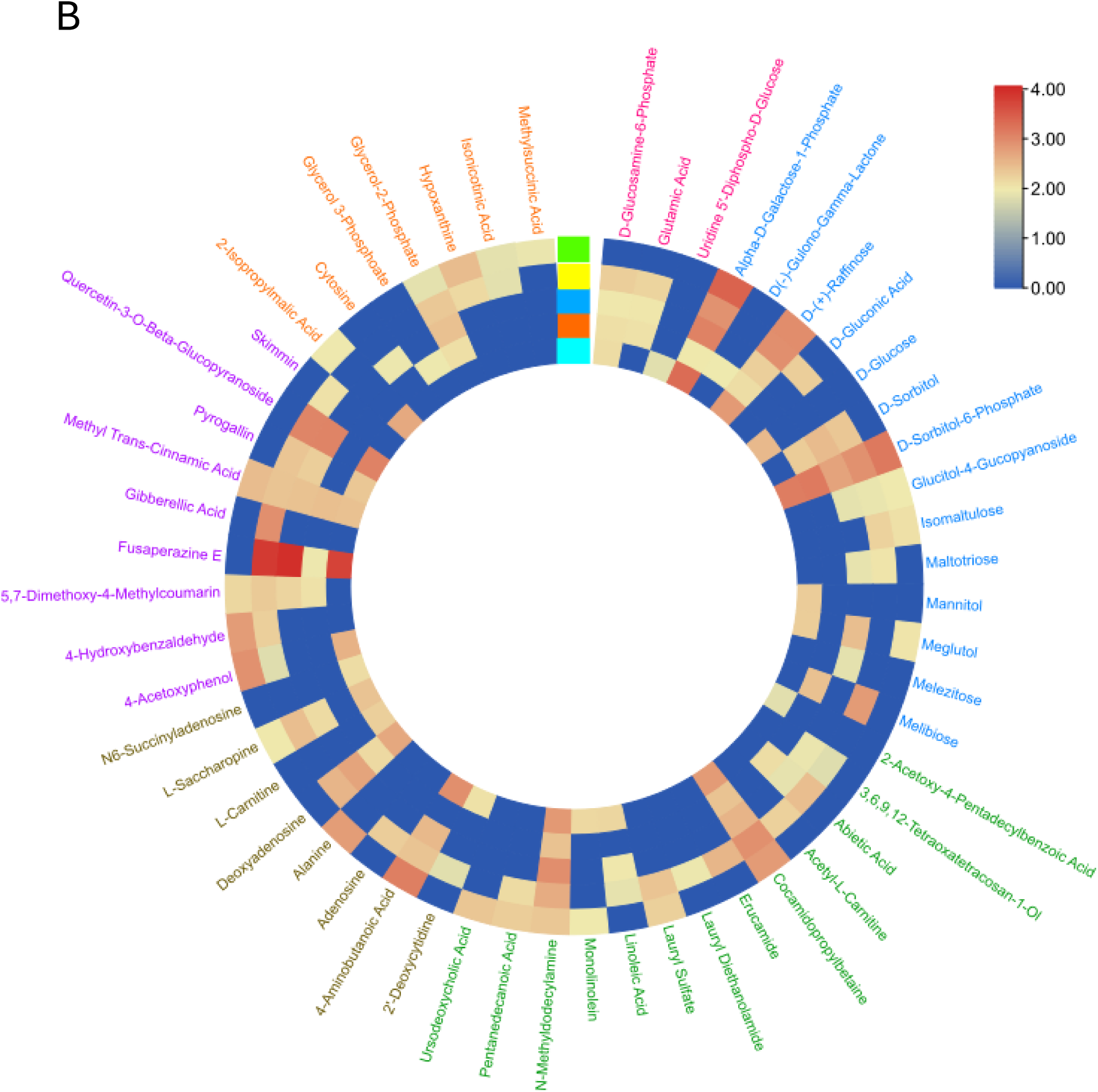
Circular heatmap of the peak area of the metabolic data. The compounds were detected in (A) fungal hyphae and (B) in the growth medium below the hyphae and cellophane. The fungi were grown as pure cultures. The metabolites found in pure media alone are subtracted in (B). The text colour indicates different chemical groups. Each block represents average values from 5 replicates.

Analysis of the secreted metabolites in the medium, below the cellophane, revealed very diverse metabolite profiles (Fig. 3B). Several compounds were secreted by all studied species, including the putatively annotated α-D*-*galactose-1-phosphate, D-raffinose, D-sorbitol, N-methyl-dodecylamine, cocamidopropyl betaine and methyltranscinnamic acid. Notably, certain compounds, including glutamic acid, abietic acid and *D*-sorbitol, were released exclusively by *T. harzianum* strains and were not detectable in *T. atrobrunneum* exudates. This indicates that the release of metabolites into the medium is both strain- and species-specific. The *L. bicolor* metabolome differed from that of *Trichoderma* strains, with compounds such as pyrolidine, thymidine, leucylglycin and histidinol not detected in *L. bicolor*, whereas they were found in all *Trichoderma* strains. Compounds such as cedrol, kaurene, *D*-sorbitol and butyl carnitine were unique only to *L. bicolor*. Categorization of metabolic features revealed 1921 and 1687 metabolites constituting the core metabolome in all *Trichoderma* strains, both in hyphae and in the medium, respectively (Fig. S4).

### Co-culture of *Laccaria* and *Trichoderma* results in strain- and species-specific global alterations of hyphal metabolomes

To understand the chemical interactions between four *Trichoderma* strains/spp. and *L. bicolor,* we analyzed the metabolome patterns of hyphae in co-culture experiments, comparing changes with PC (from Fig. 3). We considered two-time points: (i) early stage (day three) with no physical contact, i.e. information exchange possible via gas phase (in AC) and media contact (in MC) (Figs. 1, S1, S2, S3), and (ii) later stage when direct contact (in DC) was established between hyphae (day five). The results for MS8a1 and *L. bicolor* are shown in Fig. 4. Co-culture showed a similar pattern of growth inhibition as in Fig. 1. MS8a1 showed a stronger inhibitory effect on *L. bicolor* in DC (-40 ± 8%) than MC (-30 ± 5%), whereas *L. bicolor* weakly inhibited the growth of MS8a1, reducing it by 7 ± 3% in MC and 16 ± 5% in DC (Fig. 4B). Other *Trichoderma* strains (WM24a1, ES8g1, *T. atrobrunneum*) exhibited a similar behavior, with stronger growth inhibition of *L. bicolor* (Figs. S5, S6, S7).

**Fig 4.**
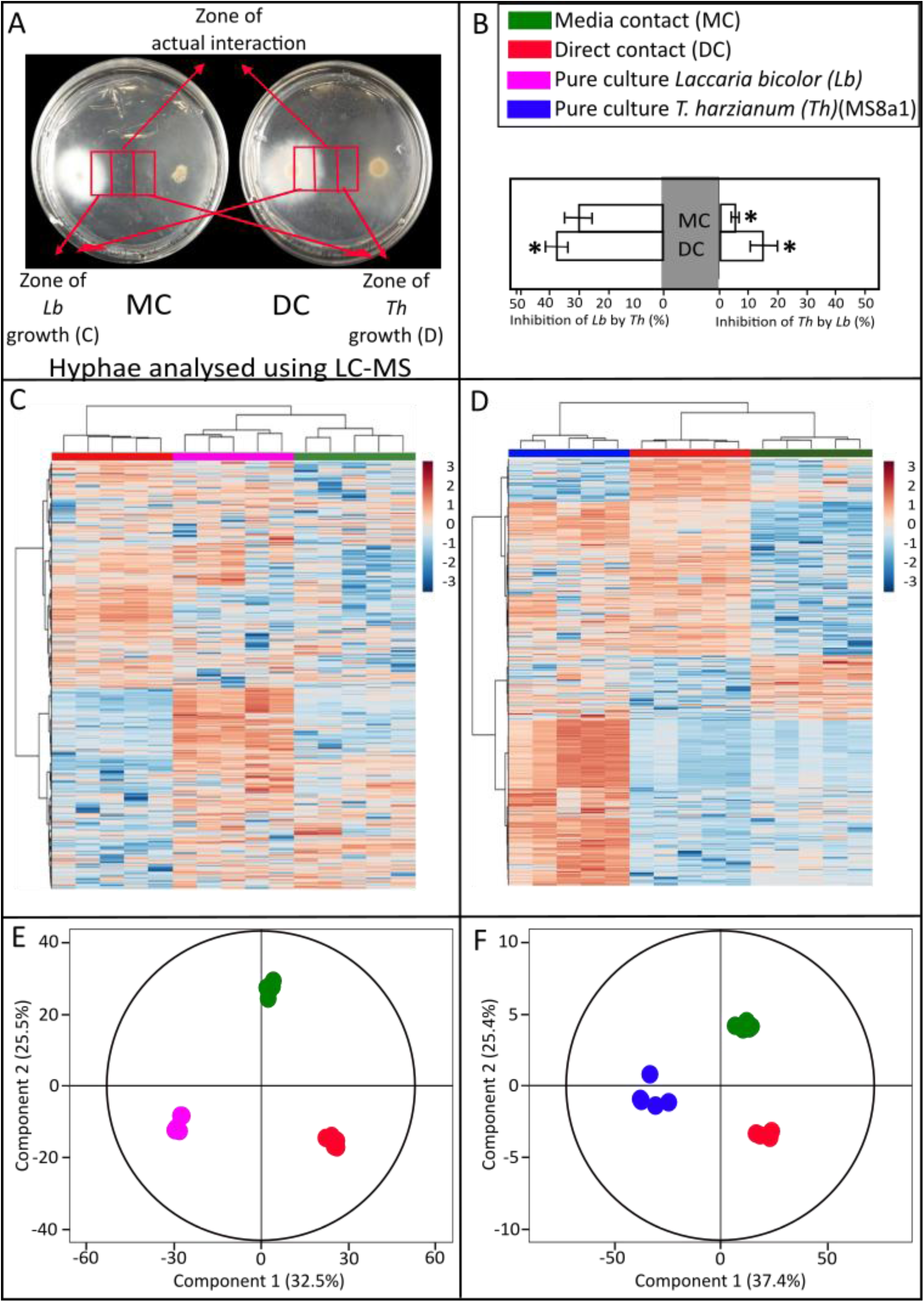
Metabolomic analysis of the hyphae from the co-cultivation experiment of *Lb* and *Th*. (A) Exemplary image of the confrontation assay showing the three different zones of sampling of hyphae and media across media contact (MC) and direct contact (DC). (B) Growth inhibition of *Th* on *Lb* (left) and vice versa (right) under different levels of co-cultivation compared to pure cultures. Significances are denoted as asterisks (one-way ANOVA and Tukey HSD, p < 0.05); mean ± SE; values are averages of 5 replicates. (C, D) Two-dimensional hierarchical clustering analysis of the peak area of features from cultures of (C) *Lb* and (D) *Th* hyphae. Features are selected based on having a Variable Importance of Projection (VIP) score >1 to compute HCA. (E, F) Orthogonal partial least square regression discriminant analysis (OPLS-DA) and adjusted p-value < 0.05 showing differences among metabolic features under different levels of co-cultivation in (E) *Lb*, OPLS model fitness: R^2^X(cum) = 0.953, R^2^Y(cum) = 1, Q^2^Y(cum) = 0.961, CV-ANOVA=1.89 x 10^-8^ and (F) *Th* OPLS model fitness: R^2^X(cum) = 0.977, R^2^Y(cum) = 1, Q^2^Y(cum) = 0.95, CV-ANOVA=4.97 x 10^-11^.

**Fig 5.**
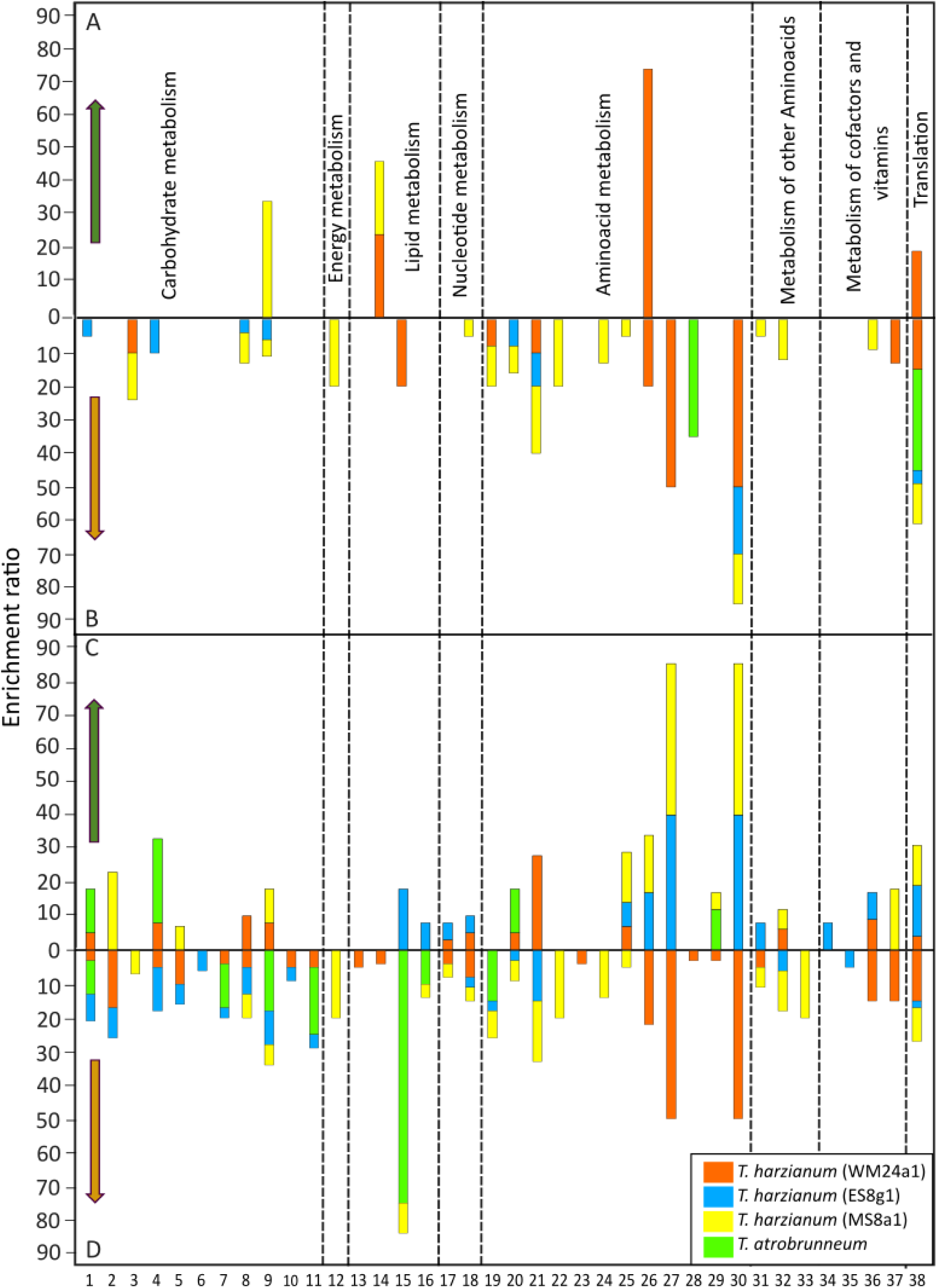
The enrichment ratios of pathways identified by significant metabolic features. The ratios are based on putative annotation in hyphae of different *Trichoderma* spp. in co-cultivation with *L. bicolor* (*Lb)* and significantly (A) up and (B) downregulated in media contact (MC) and (C) up and (D) downregulated in direct contact (DC). The numbers 1 to 38 indicate different pathways as listed in the Table 1. Pathways are grouped according to metabolic processes.

Hierarchical cluster analysis and OPLS-DA for MS8a1 (Fig. 4D & F) and *L. bicolor* (Fig. 4C & E) showed distinct metabolic patterns between PC and co-cultured mycelia. The heatmaps demonstrated global changes in the hyphal metabolomes due to interactions between MS8a1 and *L. bicolor*. The metabolic profiles allowed differentiation of scenarios along predictive component 1 (PC1; 32.5% for *L. bicolor* and 37.4% for MS8a1) and between MC and DC along PC2 (25.5% for *L. bicolor* and 25.4% for MS8a1). Similarly, the confrontation scenarios between *L. bicolor* and the other *Trichoderma* strains (Fig. S5 for WM24a1, Fig. S6 for ES8g1 and Fig. S7 for *T. atrobrunneum*) could be statistically separated based on the detected profiles, suggesting that each of the fungal strains and species developed interaction-dependent metabolic fingerprints.

Statistical analysis revealed unique responses, highlighting several metabolic features in both interacting fungi, possibly related to defence mechanisms or growth regulation. In the MC interaction between *L. bicolor* and MS8a1, 493 discriminant mass features were identified, with five upregulated and 488 downregulated compared to pure cultures. *L. bicolor* showed 423 discriminant features, of which 22 were upregulated and 401 downregulated.

In direct contact, MS8a1 hyphae had 478 discriminatory features, with 319 downregulated and 159 upregulated. *L. bicolor* had 350 discriminatory features, of which 254 were downregulated and 196 upregulated, respectively. Similar results were observed for other fungal strains. Medium contact induced upregulation of only five and six metabolic features in WM24a1 and MS8a1, respectively. *L. bicolor* showed more downregulated than upregulated features in MC with different *Trichoderma* strains. In fact, more downregulated features were detected in all co-culture conditions, except in *T. atrobrunneum*, where 79 features were downregulated and 135 upregulated in MC with *L. bicolor*.

Some of the common downregulated mass features could be putatively annotated (level 2 annotation). These include aminobutanoic acid, malic acid, valine, and triethanolamine, which appeared to be present in all *T. harzianum* strains under MC. In DC features such as L-tyrosine, uridine, lauryl sulfate, perseitol, glutathione, D-ribulose-5-phosphate, and L-arabitol were common to all *T. harzianum* strains. The total number of annotated and discriminating mass features under each co-culture condition is listed in Supplementary Table S4.

### Co-culture leads to species-specific enrichment of metabolic pathways with different functions in growth and defence

To gain a comprehensive understanding of the pathway regulation during the interaction between *Trichoderma* spp. and *L. bicolor,* we calculated the enrichment ratios of putatively annotated metabolites in the different pathways (Fig. 5). A total of 38 metabolic pathways were statistically significant in the hypergeometric test (P≤0.05) and identified as being either up- or downregulated in the hyphal cells during the antagonistic interaction between *L. bicolor* and *Trichoderma* strain/species. The changes observed in both *L. bicolor* and *Trichoderma* spp. were highly species- and strain-specific. Overall, the antagonistic interactions resulted in a generalized downregulation of pathways, which was only offset by the upregulation of a few pathways. The affected pathways are fundamental for cellular processes and growth being involved in metabolism of carbohydrates, energy, lipids, nucleotides, amino acids, cofactors, and vitamins, as well as in translation.

In the context of MC, only the *Trichoderma* strains WM24a1 and MS8a1 showed an upregulation of individual pathways. In MS8a1, metabolic pathway analysis revealed an upregulation of the glycerophospholipid and pentose pathways in response to *L. bicolor*, with enrichment ratios of 22 and 35, respectively. The greatest extent of upregulation was observed in the phenylalanine pathway in WM24a1, with an enrichment ratio of 75. Glycerophospholipid metabolism and aminoacyl-tRNA biosynthesis were also upregulated, with enrichment ratios of 25 and 20, respectively (Fig. 5A).

Conversely, downregulated, putatively annotated compounds were found to be enriched in the aminoacyl-tRNA biosynthetic pathway in MC. *T. atrobrunneum* showed an additional downregulation of the tryptophan metabolism. All three *T. harzianum* strains (WM24a1, MS8a1 and ES8g1) exhibited downregulation of compounds enriched in arginine as well as valine, leucine and isoleucine biosynthesis. In particular, *T. harzianum* (MS8a1) showed downregulation of three pathways for carbohydrates, seven for amino acids and one for each of the following pathways: nitrogen, pyrimidine, ß-alanine, glutathione, pantothenate and CoA, with varying degrees of enrichment. *T. harzianum* (WM24a1) showed downregulation in several pathways, including butanoate metabolism, linolenic acid metabolism, alanine, aspartate and glutamate metabolism, phenylalanine, tyrosine and tryptophan biosynthesis, and ubiquinone biosynthesis. *T. atrobrunneum* exhibited a reduction in the expression of four pathways involved in carbohydrate metabolism and three pathways involved in amino acid metabolism (Fig. 5B).

Upon contact (DC), a significant increase in the total number of putatively annotated compounds and enriched pathways was observed for all examined species. All *Trichoderma* spp. showed differential expression of features enriched in aminoacyl-tRNA biosynthesis. MS8a1 exhibited downregulation of three pathways related to carbohydrate metabolism, nine related to amino acid metabolism, one related to nitrogen metabolism, two related to lipid metabolism and two related to nucleotide metabolism. WM24a1 showed a reduction in the expression of eight metabolic pathways related to carbohydrate metabolism, seven pathways related to amino acid, glutathione and pantothenate metabolism, and two pathways related to lipid and nucleotide metabolism. In ES8g1, 10 pathways related to carbohydrate metabolism, three related to amino acid metabolism, one related to nucleotide metabolism and one related to pantothenate metabolism were downregulated. Compared to the *T. harzianum* strains, fewer pathways were enriched with downregulated compounds in *T. atrobrunneum*. These included four in carbohydrate metabolism, two in lipid metabolism and one pathway in amino acid metabolism (Fig. 5D).

In the DC scenario, significantly fewer upregulated pathways were detected than downregulated ones. For MS8a1, we observed an upregulation of three pathways related to carbohydrate metabolism and six each in amino acid- and ubiquinone metabolism (Fig. 5C). WM24a1 showed upregulation of four pathways involved in carbohydrate metabolism, five in amino acid, glutathione, pantothenate and CoA metabolism, whereas ES8g1 exhibited upregulation of six pathways including those involved in amino acid, pantothenate and CoA metabolism, as well as two pathways related to lipid and nucleotide metabolism. In *T. atrobrunneum*, only a subset of pathways was enriched for upregulated features, including fructose, mannose, arginine, proline and tyrosine metabolism (Fig. 5C).

### *Laccaria* and *Trichoderma* spp. hyphal exudates change during antagonism

The mycelia are in contact with the culture medium through cellophane, via which they absorb nutrients, minerals and water. The hyphae in turn secrete metabolites into the medium (Fig. 4). Thus, in addition to the gas phase (Figs. 1, 2), the medium is one of the ways in which the two competitors can communicate. To gain further insight into the complex chemistry involved in the interaction between the two plant beneficial fungi, we developed an experimental setup to analyze the metabolic composition of the chemical exudation (Fig. 6A). We analyzed culture media segments i) directly beneath the hyphae ii) in the interphase between the fungi before direct physical contact (MC) and later iii) directly below the hyphae of the two fungi in DC with each other. After removal of the cellophane between the hyphae and media, the metabolic exudates from the three zones were analyzed (Fig. 6B). The HCA and OPLS-DA analyses revealed different metabolite patterns depending on the degree of interaction between *Laccaria* and MS8a1 (Fig. 6C, E). Similarly, clear pattern differences in the metabolic profiles were observed for the other *T. harzianum* strains (Fig. S8 and S9), and for *T. atrobrunneum* (Fig. S10). *L. bicolor* also responded individually to the presence of each of the *Trichoderma* species, as evidenced by changes in the metabolite secretion (Fig. 6 D, F). The observed changes in the metabolite pattern of *L. bicolor* were dependent on the partner strain/species encountered, suggesting a specific, strain-dependent exudation response in the ECM fungus (Figs.6, S8, S9, S10 (D, F)).

**Fig 6.**
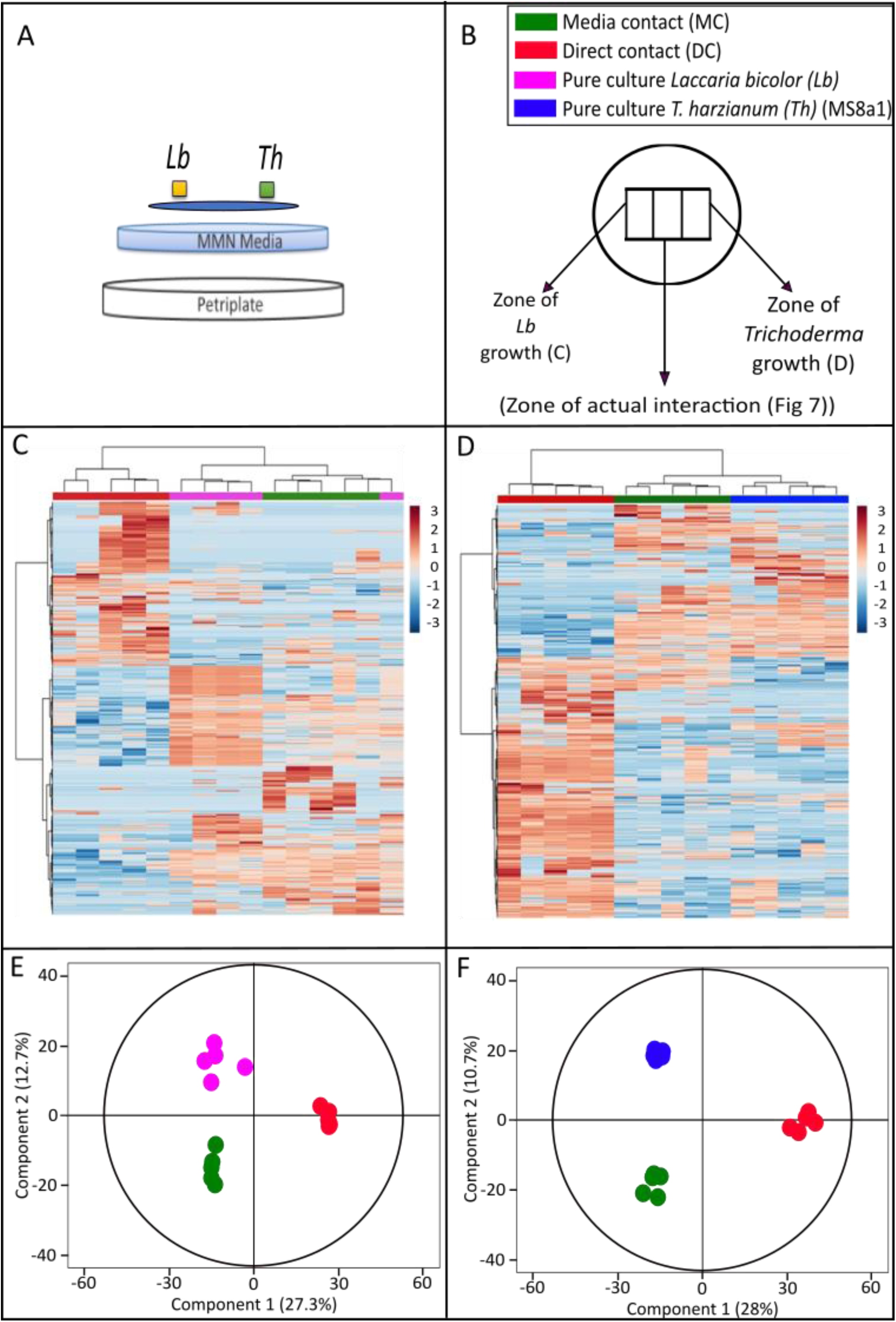
Metabolomic analysis of the media from the co-cultivation experiment of *Lb* and *Th.* (A) Schematic representation of the co-cultivation experiment between *Lb* and *Th* showing the position of fungal inoculants from the two species, cellophane sheet, and the underlying media. (B) Experimental set-up of the confrontation assay showing the three different zones of sampling of media across media contact (MC) and direct contact (DC). (C,D) Two-dimensional hierarchical clustering analysis of the peak area of features from cultures of (C) *Lb* and (D) *Th.* (E, F) Orthogonal partial least square regression discriminant analysis (OPLS-DA) showing differences among metabolic features in media under different levels of co-cultivation in (E) *Lb*, OPLS model fitness: R^2^X(cum) = 0.953, R^2^Y(cum) = 1, Q^2^Y(cum) = 0.799, CV-ANOVA=6.45 x 10^-4^ and (F)*Th*, OPLS model fitness: R^2^X(cum) = 0.98.7, R^2^Y(cum) = 1, Q^2^Y(cum) = 0.785, CV-ANOVA=4.23 x 10^-4^.

All the heatmaps based on the *Laccaria*-*Trichoderma spp*. interaction metabolites showed distinct patterns in MC (Figs. 6C & D, and Figs. S8, S9, S10). The OPLS-DA demonstrated a clear separation of the metabolic profiles between the exudates of the pure cultures and the co-culture scenarios along PC2 (12.7 % for *L. bicolor* and 10.7 % for MS8a1 and between MC and DC scenarios along PC2 (27.3 % for *L. bicolor* and 28 % for MS8a1 (Fig. 6E & F). Clear separation in the OPLS-DA was also evident in the confrontations of *L. bicolor* with *T. harzianum* (WM24a1 and ES8g1) and with *T. atrobrunneum* (Figs. S8, S9, S10).

Of the 92 discriminant features in the medium under the mycelium of MS8a1, 22 were upregulated and 70 were downregulated. In *L. bicolor* samples in contact with MS8a1 only 78 discriminating metabolites were detected in the exudate with 19 metabolites upregulated and 59 downregulated.

Direct contact resulted in 159 discriminating mass features under the MS8a1 mycelium, with 47 features downregulated and 112 features upregulated. Under *L. bicolor* mycelium, only 15 downregulated features were detected, which may be due to the overgrowth by MS8a1 hyphae. The total number of discriminatory features under each co-culture condition for each fungus examined is shown in Supplementary Table S5. While not many differentially expressed features could be annotated, the highest upregulation of features was observed under MS8a1 mycelium in DC with *L. bicolor*. Compared to the other *Trichoderma* strains*, T. atrobrunneum* exuded fewer active metabolites with only four and three features upregulated in DC and MC, respectively. Fewer differentially regulated compounds were also detected from *L. bicolor* in DC than in other co-culture scenarios, suggesting that *L. bicolor* released generally fewer compounds towards the end of the co-culture with *Trichoderma* spp.

### Exudate composition depends on the distance between the interacting fungi

The composition of diffusible metabolites included nucleotides, amino sugars, carbohydrates, peptides and amino acids, lipids and phytochemical oxyaromatic compounds for all the fungi analyzed at different degrees of contact (Fig. 7). An alluvial plot allowed analysis of the flow and diffusion of exudates from each fungus into the contact zone. The analysis revealed six distinct, species-dependent patterns of mass flow for all *Laccaria* - *Trichoderma* interactions (Fig. 7A). Overall, the distinguishing mass features in MS8a1 exudates were predominantly oxyaromatic compounds, followed by lipids, peptides and carbohydrates. In all other *T. harzianum* strains (WM24a1, ES8g1 and *T. atrobrunneum*), the predominant secreted compounds were lipids, followed by oxyaromatic compounds, peptides and carbohydrates (Fig. 7C). Depending on the mass feature, the concentration either decreased in the interaction zone or, feature was completely novel and detected only in the interaction zone compared to the other zones studied.For example, in the case of *L. bicolor* and MS8a1, many of the mass features that were detected directly below the fungus were not found in the interaction zone (Fig. 7A). The analysis of the exuded metabolites of *T. harzianum* (WM24a1, ES8g1) and *T. atrobrunneum* in interaction with *L. bicolor* showed similar results: the composition of the detected mass features was different directly below the fungus compared to the interaction zone (Fig. S11). In addition to the species-specific exudation patterns in *Trichoderma* spp, we also observed differences in the exudation of *L. bicolor* depending on the interaction partner (Fig. 7A; Fig. S11).

**Fig 7.**
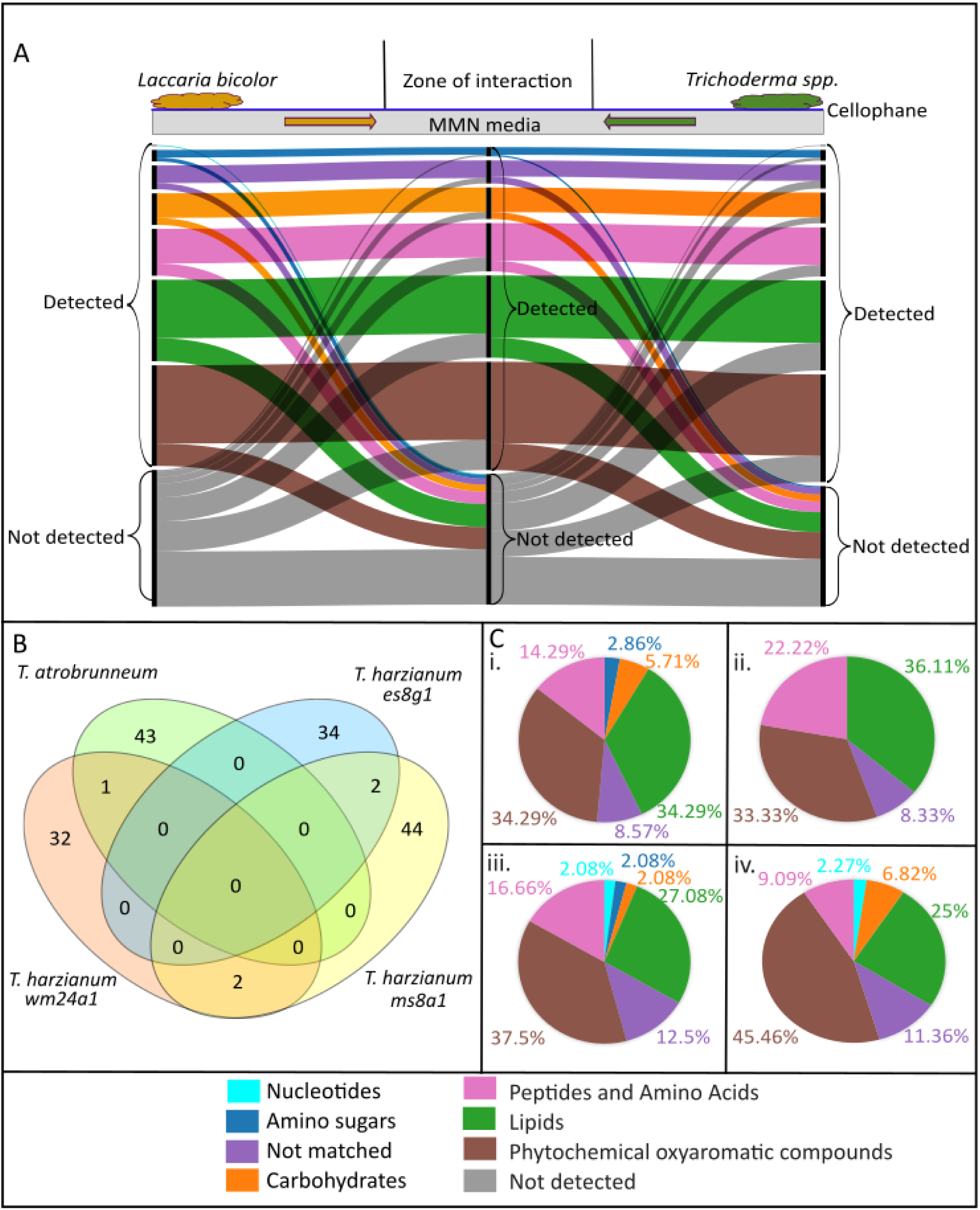
Metabolomic analysis of the media from the zone of actual interaction between *Lb* and *Th.* (A) Representative alluvial plot showing the flow of exudates from *L. bicolor* to *T. harzianum (*MS8a1) and *vice versa* through the common growth media in co-cultivation (read from left to right). The alluvial plots for other species are shown in Fig. S11.The colour patterns depict features belonging to different chemical groups classified by ‘multidimensional stoichiometric compound classification’ (MSCC). (B) Venn plot depicting the exudates found ‘uniquely’ in the zone of interaction across four different species. (C) Chemical class classification of the compounds by MSCC of the features found ‘only’ in the zone of interaction across co-cultivation of (i) *T. harzianum (*WM24a1), (ii) *T. atrobrunneum*, (iii) *T. harzianum (*ES8g1) and (iv) *T. harzianum* (MS8a1) with *L. bicolor*.

Concerning the detected differences in the mass feature patterns, the accumulation of exclusively in the interaction zone compared to PC is particularly noteworthy. The composition of accumulated metabolites in the interaction zone differed between *L. bicolor* and the four *Trichoderma* confrontation experiments. All three *T. harzianum* strains (WM24a1, MS8a1 and ES8g1) and *T. atrobrunneum* showed unique metabolic mass characteristics in the interaction zone, with 32, 44, 34 and 43 compounds, respectively (Fig. 7b). WM24a1 and MS8a1, as well as MS8a1 and ES8g1, each shared two common mass features. In contrast, only one common mass feature was detected between WM24a1 and *T. atrobrunneum* (Fig. 7b). However, these common compounds could not be annotated.

Some of the mass features detected in the interaction zone could be annotated as phenolic compounds with antifungal properties. These compounds included quercetin 3-O-rutinoside in *T. harzianum* (WM24a1) interaction zone, salicylic acid and pyrogallin in *T. harzianum* (ES8g1) interaction zone, and vanillin in *T. harzianum* (MS8a1) interaction zone. We detected moreover some metabolites that were specific for *T. harzianum* strains and were not detected in *T. atrobrunneum* interactions. Such compounds detected exclusively in the interaction zone of *L. bicolor* - *T. harzianum* strains included minimal amounts of nucleotides, amino sugars, and carbohydrates.

## Discussion

### Restricted mycelial growth and altered VOC profiles in *L. bicolor* – *Trichoderma* co-cultures

Due to their volatility and diffusivity, VOCs are the ideal signalling molecules between interacting fungi even before direct hyphal contact (Schulz-Bohm et al., 2017). To function as signalling molecules, unique, adaptable VOC profiles are considered as a prerequisite (Rosenkranz et al., 2021). Our study reveals highly unique VOC profiles under solitary cultures but also under dual cultivation for all species and interaction scenarios investigated. The profiles were, furthermore, adapted depending on the degree of contact with other fungal species, thus suggesting function in fungal interaction and supporting our first hypothesis (I). Unique VOC profiles were previously detected also from different *T. harzianum* strains in pure cultures (Lee et al., 2019; Nemčovič et al., 2008; Nieto-Jacobo et al., 2017; Siddiquee et al., 2012,2014). For example, Lee et al. (2019) detected 27 VOCs from *T. harzianum* CBS 227.95 and Siddiquee et al. (2012) even 278 VOCs from the *T. harzianum* strain FA1132. Moreover, when three other *Trichoderma* species (*T. hamatum* QL15d1, *T. reesei* QM6a and *T. velutinum* GL1561) were studied as pure cultures and in co-culture with *L. bicolor,* highly unique, species and interaction specific VOC profiles were reported (Guo et al., 2019).

In accordance with Guo et al. (2019), the present results show that the interaction of *Trichoderma* spp. with *L. bicolor* leads to strong interaction-dependent adjustments in the VOC profiles even if headspace VOC concentrations tended to be reduced in dual cultures compared to PC. An exception was the WM24a1 - *L. bicolor* interaction in AC, which also appears to be special in other respects: Transcriptomic data from Stange et al. (2024) showed that in a similar co-culture set-up in which the communication was allowed through VOCs, KEGG pathways associated with the biosynthesis of secondary metabolites were upregulated in *L. bicolor.* The authors observed, moreover, a downregulation of WM24a1 terpene synthase gene (M431DRAFT_113113). In line with these results, in the present study, *L. bicolor* growth was overall only marginally affected in AC with *Trichoderma* and especially WM24a1, suggesting that the *Trichoderma* VOCs have no or only a weak effect on *L. bicolor* growth, whereas ECM VOCs may negatively affect *Trichoderma* performance. At later stages, however, when also exchange of soluble metabolites were allowed, *Trichoderma* behaved more aggressively by overgrowing *L. bicolor* under *in vitro* conditions. Somewhat contradictory to the growth measurements, that revealed inhibition of *Trichoderma* growth in the presence of *Laccaria* VOCs, in general reduced *L. bicolor* VOC profiles were observed in co-cultures. The results suggest that either the ECM compounds are somehow degraded in the presence of *Trichoderma* spp. or, alternatively, *L. bicolor* may simply release other VOCs or invests more in other, soluble metabolites instead of VOCs when sensing a competitor. Together the present results of fungus-specific and interaction degree-dependent changes in VOC profiles strongly support the hypothesis that fungi can regulate their VOC emissions in response to environmental constraints. This adjustability suggests that fungal VOCs possess important ecological functions in microbial interactions and perception.

### Altered metabolic responses in the hyphae and exudates of *Trichoderma* and *L. bicolor* in co-cultures

Analyses of core metabolomes are essential to understand the fungal responses to various environmental changes. The present metabolite analyses highlight pronounced similarities among *T. harzianum* strains and species-specific metabolic diversity with *T. atrobrunneum*. A comprehensive metabolite analysis of hyphae and exudates in pure cultures identified 9365 mass features with varying levels of putative annotation (Level 2-5). This limits the exploration of individual metabolites but allows the classification of functional chemical composition. Our hypothesis (II), that there is a high degree of metabolic similarity within the studied *Trichoderma* spp., is however, only weakly supported by the data: While similar carbohydrate composition was detected in the hyphae of the different *T. harzianum* strains, there were notable strain-dependent differences in lipids, peptides, and phytochemicals. Secreted metabolite profiles also varied: Compounds such as α-*D*-galactose-1-phosphate and *D*-raffinose were common to all studied fungi whereas, in the hyphae of the studied ECM unique metabolites such as cedrol, kaurene, *D*-sorbitol, and butyl carnitine were detected. The core metabolomes can serve as reference metabolome to identify the changes in hyphal and exudate composition that may facilitate chemical exchange between interacting fungi (Mukherjee et al., 2012).

*Trichoderma* species are known to produce exudates that degrade microbial cells in soil habitats, altering the ability of other species to absorb nutrients and persist (Assigbetsé et al., 2012). *T. harzianum, for example,* can alter pathogenic fungal communities and some of its metabolites (incl. harzianopyridone, pyrone, and trichorzianine) show antifungal activity (Khan et al., 2020; Umadevi et al., 2018; Vinale et al., 2008). How *Trichoderma* spp. interacts with non-plant pathogenic fungi is less well understood. Our study reveals for the first-time insight to the soluble metabolites potentially involved in interactions between *Trichoderma* spp. and an ECM. The results show that co-culture with the *L. bicolor* leads to distinct strain- and species-specific changes in the exudates and in the hyphal metabolites of *Trichoderma*. These alterations include global changes that distinguish pure cultures from co-culture scenarios (MC and DC) in all studied *Trichoderma* strains, suggesting interaction-dependent metabolic fingerprints. The altered metabolome may have also directly influenced the fungal growth: *Trichoderma* constantly inhibited *L. bicolor* growth more than *vice versa* when the exchange of soluble metabolites was allowed.

Some common features were detected in the hyphal metabolome of the different *T. harzianum* strains, including the upregulation of such compounds as L-tyrosine, uridine, lauryl sulfate, perseitol, glutathione, D-ribulose-5-phosphate, and L-arabitol in DC. In MC all the studied *T. harzianum* strains showed downregulation of aminobutanoic acid, malic acid, valine, and triethanolamine. These results highlight that the metabolic responses that occur upon interaction with other fungi are at least to some extent shared among the different strains. However, whereas the metabolites detected within the hyphae are interesting allowing direct insight to the altered fungal metabolism, the metabolites secreted into the growth matrix are likely to play a more critical role in fungal interactions. Statistical analysis revealed many differentially regulated metabolic features in the *Trichoderma* exudates. Many of these were previously assigned a role in fungal defence or inhibition (discussed below). The differential regulation suggests specific chemical interactions and activated defence mechanisms upon sensing a competitor. Striking were also the notable differences detected in the exudate profiles of *L. bicolor* confronted with different *Trichoderma* spp. These findings suggest that the ECM can sense the competitor and show a unique exudation response towards a specific *Trichoderma* strain. The results challenge the hypothesis that there would be a conserved exudation response in *L. bicolor* towards different antagonistic fungi.

### Co-cultivation led to altered metabolic pathway enrichment influencing growth and defence during fungal interactions

As the fungi grow towards each other, they continuously sense their environment and initiate specific downstream pathways based on the prevailing abiotic and biotic conditions (Bahn et al., 2007; Hinterdobler et al., 2021; Sarma et al., 2014; Turrà & Di Pietro, 2015; Zeilinger & Atanasova, 2020). Enrichment analysis of the putatively annotated metabolic features (level 2) revealed differential regulation of several metabolic pathways involved in carbohydrate, energy, lipid, nucleotide and amino acid metabolism. Altogether, we identified 38 metabolic pathways that were altered during the antagonistic interactions, although downregulation was observed more frequently than upregulation. For example, all *T. harzianum* strains exhibited a reduction in pathways associated with arginine and valine, leucine, and isoleucine biosynthesis on MC. These alterations together with observed reduced growth may reflect the increased investment to defence or interaction related metabolites and less to growth. In general, such response of *T. harzianum* strains was stronger compared to *T. atrobrunneum*, former showing more metabolic pathways enriched by downregulated compounds.

Upregulated pathways in the DC scenario were significantly fewer than downregulated ones. A few commonalities among the upregulated pathways were found especially among the *T. harzianum* strains, though. For example, MS8a1, WM24a1 and ES8g1 all showed upregulation in pathways related to amino acid metabolism, whereas in MS8a1 and WM24a1 carbohydrate metabolism related pathways were altered. Ubiquinone metabolism related changes were, however, specific to MS8a1 and pantothenate and CoA metabolism related upregulation specific to ES8g1. In *T. atrobrunneum,* only a subset of pathways was enriched for upregulated features eventually reflecting less aggressive defence reaction compared to *T. harzianum* strains in the presence of *L. bicolor*,

Together the detected alteration in various pathways shows that the interacting fungi can sense each other through signalling compounds thus supporting our hypothesis (III). The results are also in accordance with the recent findings of Stange et al. (2024), who analyzed the transcriptional changes of *L. bicolor* and *T. harzianum* WM24a1 in co-culture scenarios and revealed an increase in the number of differentially expressed genes (DEGs) in DC compared to MC. The authors revealed DEGs mainly assigned to KEGG metabolic pathways involved in the biosynthesis of secondary metabolites, possibly indicating the activation of secondary metabolism-based communication, especially related to linolenic acid metabolism and leucine and isoleucine degradation (Stange et al., 2024). Interestingly, also the present analyses revealed that linolenic acid metabolism was downregulated in WM24a1 under MC. Valine, leucine and isoleucine biosynthesis were, moreover, downregulated under both MC and DC. Together with the transcriptional analyses (Stange et al., 2024), the present metabolomic analyses serve as a first step towards understanding the complex interactions and molecular regulation behind the fungal interaction and support the hypothesis that fungi alter their metabolic pathways in response to their biotic environment.

### Co-culture led to varying metabolic exudates with a potential role in communication at the confrontation zone in MC

In the soil matrix as well as in *ex-situ* co-culture experiments, the diffusion capability of allelochemicals through the matrix play a crucial role in fungal interactions (Shahriar et al., 2022). The changes in the composition and distribution of the exudates over distance is likely to alter the interaction result and defence. Using alluvial plots, we analysed the flow and diffusion of exudates from *L. bicolor* and *Trichoderma* mycelia until the contact zone and detected six distinct patterns of mass flow in the studied interactions. These results provide clear examples of the potential metabolic changes during *L. bicolor* - *Trichoderma* confrontation. Interesting are the changes over distance, reflected as degraded as well as novel compounds detected in the contact zone compared to PC. The specific composition of these metabolites was dependent on the interaction partner, highlighting species-specific recognition and antagonism. In line with other present results, these analyses revealed *Trichoderma* strain-specific differences in the response of the ECM.

Some common mass features were detected from all *Trichoderma* strains; however, annotation was possible only for the discriminating features that included mainly oxyaromatic compounds, lipids and peptides. When *L. bicolor* was co-cultured with any of the *T. harzianum* strains, nucleotides, amino sugars, and carbohydrates were detected in the interaction zone, while these compound classes were not dominant in co-culture with *T. atrobrunneum.* Interestingly, some potentially fungal defence related phenolic compounds, such as quercetin 3-O-rutinoside, salicylic acid, pyrogallin and vannilin, were identified in the interaction zone between *L. bicolor* and *Trichoderma spp.* These have been previously reported to inhibit the growth of *T. harzianum* (Gómez-Vásquez et al., 2004, Reino et al., 2008; Philip et al., 2024) and might also be a cause of the growth inhibition we detected in the co-cultures. Altogether these diffusion patterns support the hypothesis (IV) that fungi could release signalling metabolites into the growth matrix long before the hyphae can encounter each other (MC), which helps to avoid/ defend a competing fungal species.

## Conclusions

In this study, we combined volatile and soluble metabolomic data to investigate the chemical processes underlying interactions between plant-beneficial fungi that might exhibit antagonistic behaviour. Notwithstanding the restricted power of *in vitro* co-cultures in the laboratory and the limited annotation of metabolites, our study offers compelling evidence for a high degree of metabolic diversity in the interacting fungi. The data illustrate that both plant-beneficial fungal groups are capable of recognizing the chemical differences between their respective interaction partners and subsequently reorganising their metabolism in response. It must be acknowledged that our study cannot replace field studies. However, it does provide a foundation upon which further investigation of analogue interactions under more natural conditions and three-partite interactions with the plant host can be conducted. In conclusion, the results of this study demonstrate the potential of volatile and soluble allelochemicals in mediating interspecific communication, recognition and competition between *Trichoderma* and an ECM (*L. bicolor*).

## List of abbreviations

Th: Trichoderma harzianum
T.arb: Trichoderma atrobrunneum
Lb: Laccaria bicolor
AC: Aerial Contact
MC: Media Contact
DC: Direct Contact
VOC: Volatile Organic Compound
MT: Monoterpenes
SQT: Sesquiterpenes
oMT: oxygenated Monoterpenes
oSQT: oxygenated sesquiterpenes
PCA: Principal Component Analysis
OPLS-DA: orthogonal Partial Least Square regression -Discriminant Analysis
HCA: Hierarchical Clustering Analysis ()

## Declarations

### Ethics approval and consent to participate

**-**

### Adherence to national and international regulations

**-**

### Consent for publication

**-**

### Availability of data and materials

The datasets generated and/or analysed during the current study are available in the open science framework data repository and can be accessed following the link: https://osf.io/c357a/?view_only=7397fa80cc9846869879168112478139

### Author contributions

Conception and design: Jörg-Peter Schnitzler, Maaria Rosenkranz, J. Philipp Benz and Prasath Balaji Sivaprakasam Padmanaban. Experimentation, Data acquisition, analysis and visualization: Prasath Balaji Sivaprakasam Padmanaban, Pia Stange, Jörg-Peter Schnitzler, Maaria Rosenkranz, J. Philipp Benz, Andrea Ghirardo and Karin Pritsch. Original draft preparation: Prasath Balaji Sivaprakasam Padmanaban. Review and editing: Maaria Rosenkranz, Jörg-Peter Schnitzler, Andrea Ghirardo, J. Philipp Benz, Karin Pritsch, Pia Stange and Tanja Karl. Project administration: Maaria Rosenkranz, J. Philipp Benz and Jörg-Peter Schnitzler.

## Acknowledgements

The authors thank Michael Witting (Helmholtz Munich, Metabolomics and Proteomics Core Facility) for valuable advice on data analysis, Ina Zimmer (Helmholtz Munich, Research Unit Environmental Simulation) for valuable advice and help with maintaining the fungal cultures and Monika Schmoll (AIT Austrian Institute of Technology GmbH, Austria) for the gift of the used *Trichoderma harzianum* strains.

## Supplementary information

Supplementary Material 1.

