## Supplementary material for "Deciphering plant-beneficial fungal interactions: Unravelling metabolic diversity that underpins communication between *Laccaria bicolor* and *Trichoderma*"

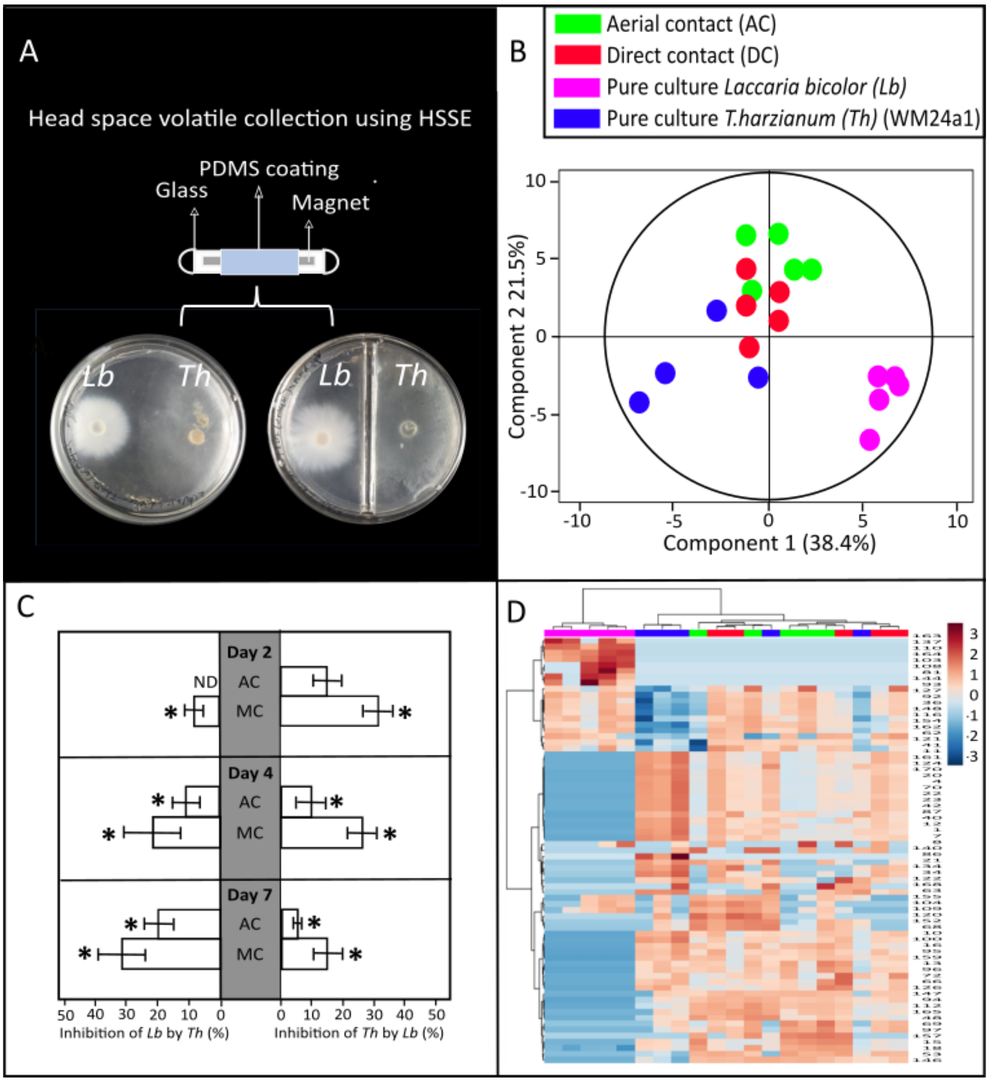

**Fig S1.** VOC analysis of of *Laccaria bicolor (Lb)* and *T. harzianum (Th)* (WM24a1*) (Th)* co-cultivated either in aerial contact (AC) or media contact (MC). (A) Experimental setup of the VOC collection from the fungal AC and DC co-cultivation in split and non-split petri plates by twisters employing Headspace sorptive extraction (HSSE) technique. (B) orthogonal partial least square regression discriminant analysis (OPLS-DA) showing differences among VOC profiles under different levels of co-cultivation, (C) Growth inhibition of *Th* on *Lb* (Left) and *Lb* on *Th* (Right) under different levels of co-cultivation compared to growth on pure cultures. ND: not detected. Significances within each day are denoted as asterisks (one-way ANOVA and Tukey HSD, $p < 0.05$); mean ± SE; Values are average of 5 replicates and (D) Hierarchical clustering analysis of the volatile concentration of the VOCs from the two fungi grown as MC, AC or pure culture*.*

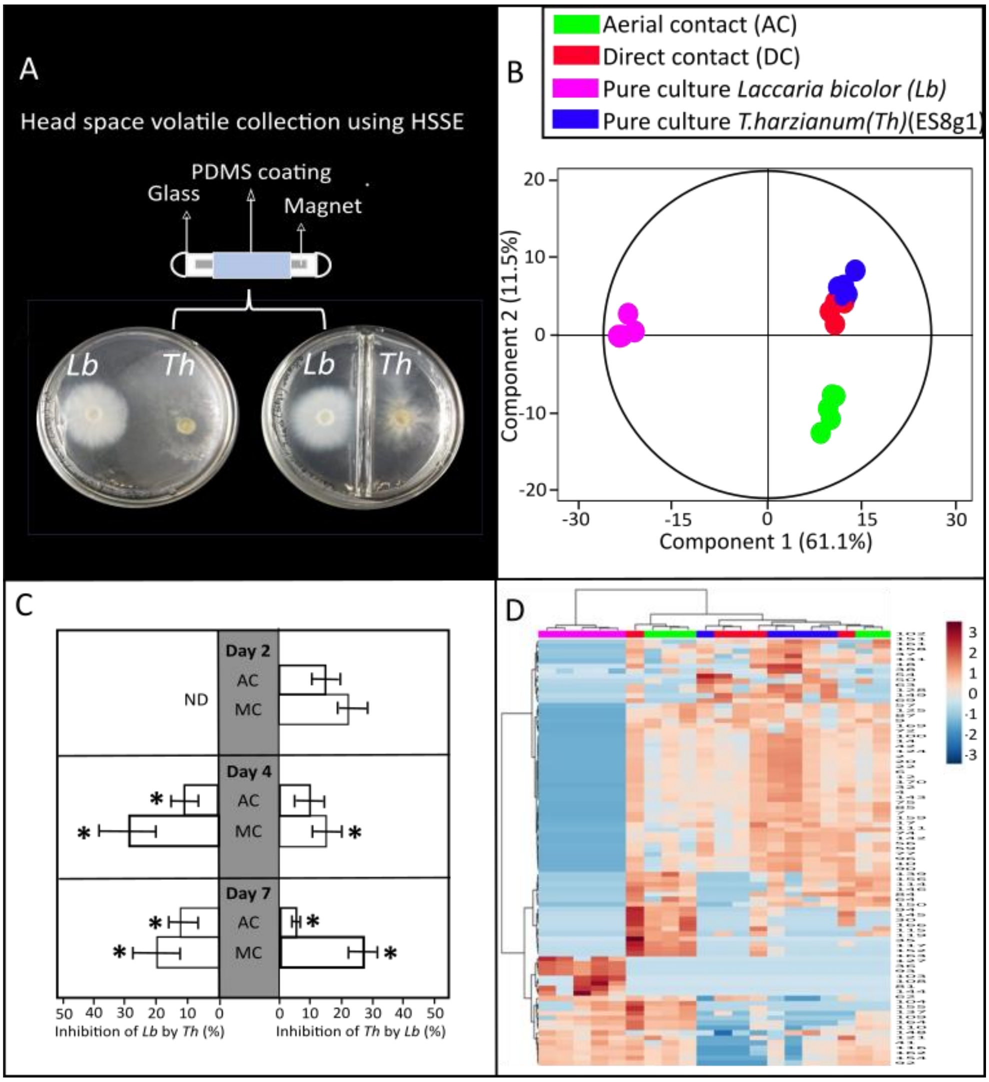

**Fig S2.** VOC analysis of of *Laccaria bicolor (Lb)* and *T. harzianum (Th)* (ES8g1) *(Th)* co-cultivated either in aerial contact (AC) or media contact (MC). (A) Experimental setup of the VOC collection from the fungal AC and DC co-cultivation in split and non-split petri plates by twisters employing Headspace sorptive extraction (HSSE) technique. (B) orthogonal partial least square regression discriminant analysis (OPLS-DA) showing differences among VOC profiles under different levels of co-cultivation, (C) Growth inhibition of *Th* on *Lb* (Left) and *Lb* on *Th* (Right) under different levels of co-cultivation compared to growth on pure cultures. ND: not detected. Significances within each day are denoted as asterisks (one-way ANOVA and Tukey HSD, $p < 0.05$); mean ± SE; Values are average of 5 replicates and (D) Hierarchical clustering analysis of the volatile concentration of the VOCs from the two fungi grown as MC, AC or pure culture.

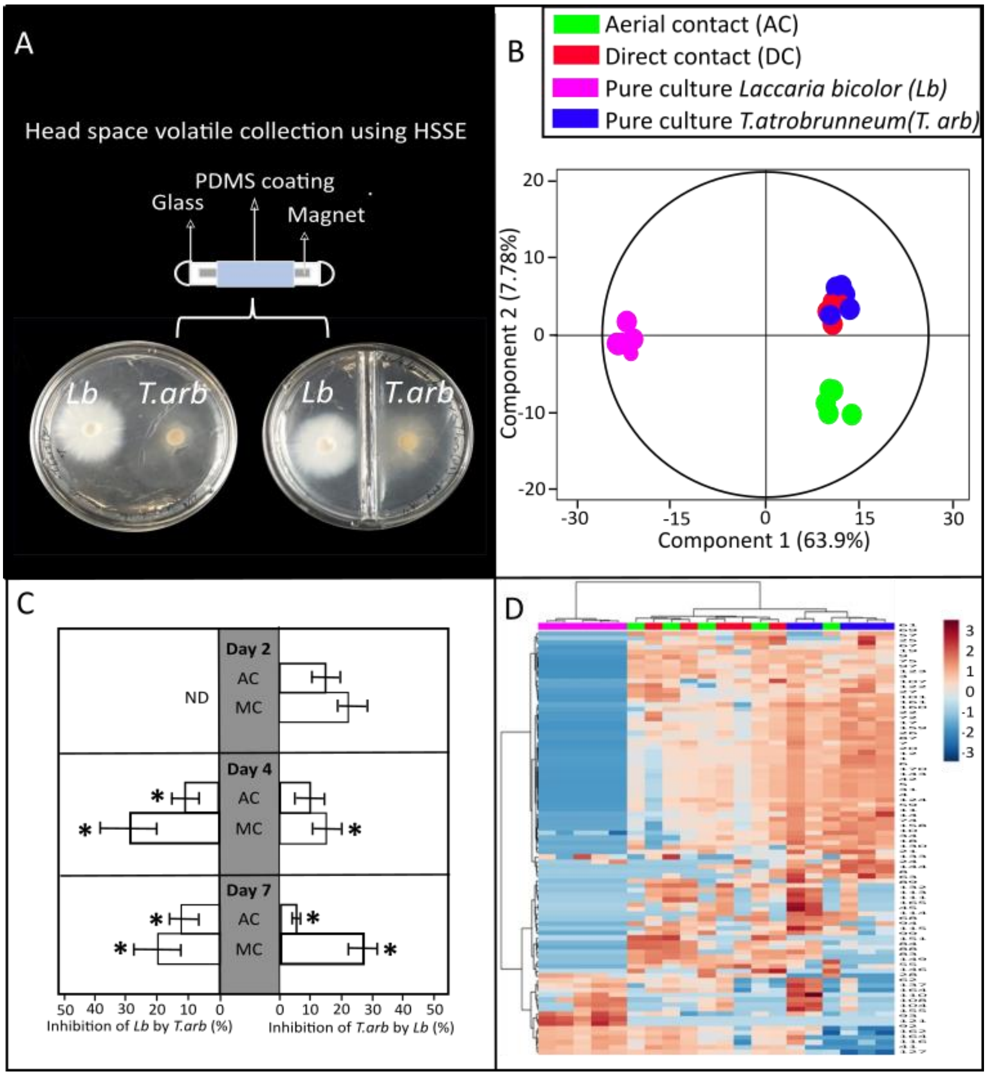

**Fig S3.** VOC analysis of of *Laccaria bicolor (Lb)* and *T. atrobrunneum (T.arb)* co-cultivated either in aerial contact (AC) or media contact (MC). (A) Experimental setup of the VOC collection from the fungal AC and DC co-cultivation in split and non-split petri plates by twisters employing Headspace sorptive extraction (HSSE) technique. (B) orthogonal partial least square regression discriminant analysis (OPLS-DA) showing differences among VOC profiles under different levels of co-cultivation, (C) Growth inhibition of *T.arb* on *Lb* (Left) and *Lb* on *T.arb* (Right) under different levels of co-cultivation compared to growth on pure cultures. ND: not detected. Significances within each day are denoted as asterisks (one-way ANOVA and Tukey HSD, $p < 0.05$); mean ± SE; Values are average of 5 replicates and (D) Hierarchical clustering analysis of the volatile concentration of the VOCs from the two fungi grown as MC, AC or pure culture.

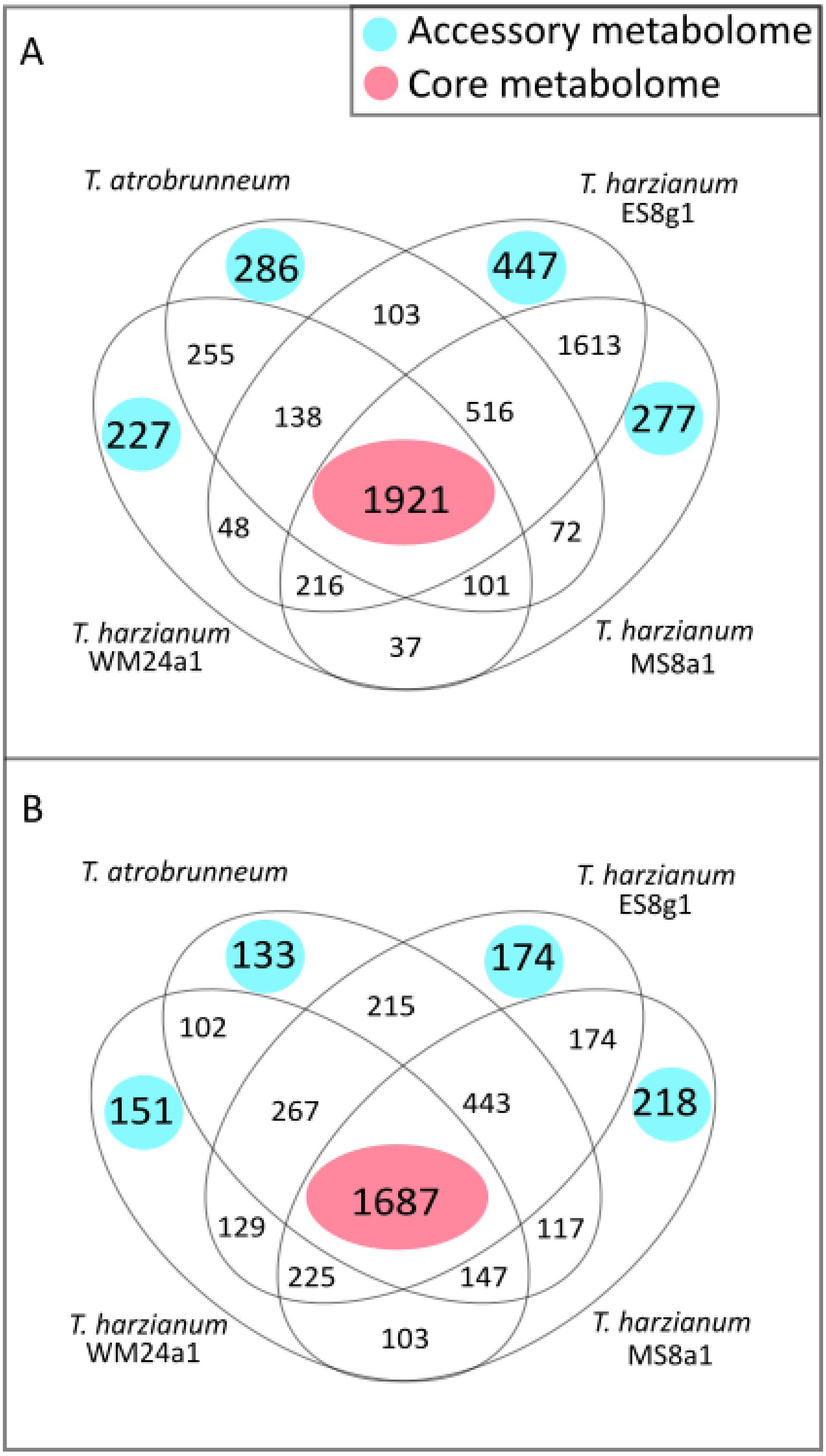

**Fig S4.** Accessory (unique) and core (common) metabolome in the hyphae and media of pure cultures in different *Trichoderma* species.

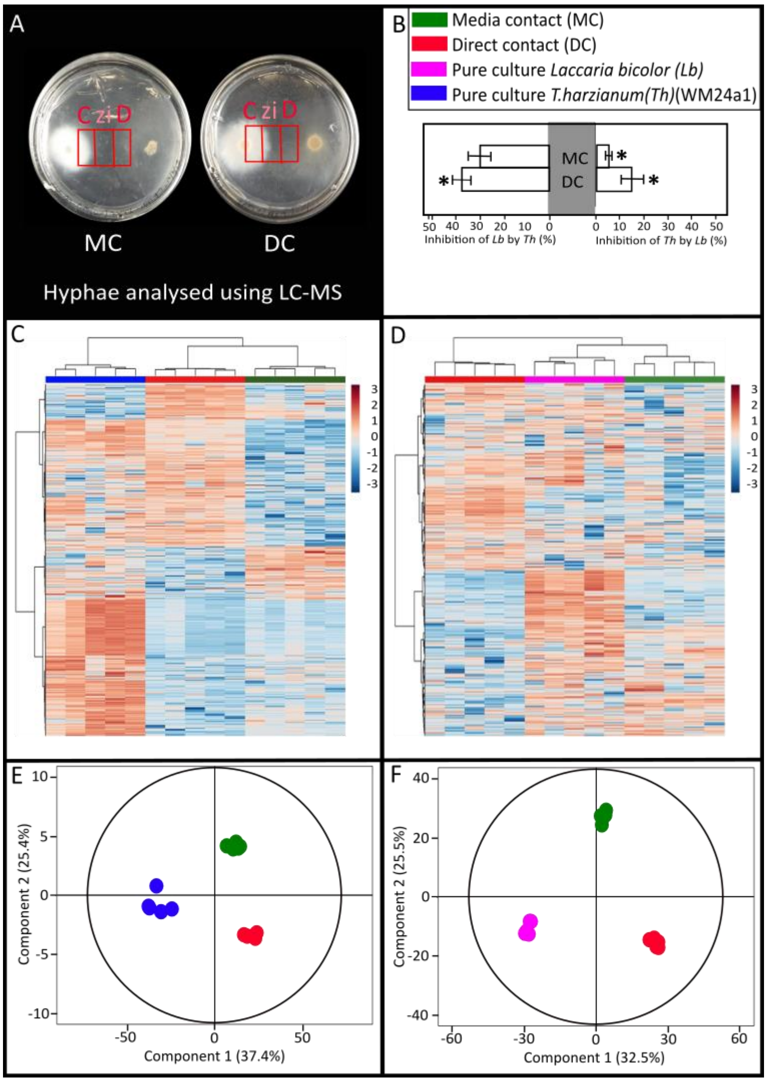

**Fig S5.** Metabolomic analysis of the hyphae from the co-cultivation experiment of *Laccaria bicolor* (*Lb*) and *T. harzianum (Th)* (WM24a1*).* (A) Exemplary image of the confrontation assay showing the three different zones of sampling of hyphae and media across media contact (MC) and direct contact (DC). (B) Growth inhibition of *(Th)* on *Lb* (left) and vice versa (right) under different levels of co-cultivation compared to pure cultures. Significances are denoted as asterisks (one-way ANOVA and Tukey HSD, $p < 0.05$); mean ± SE; Values are average of 5 replicates. Hierarchical clustering analysis of the peak area of features from cultures of (C) *Lb* and (D) *(Th)* hyphae grown in media contact (MC) direct contact (DC) and pure cultures. Features are selected based on having a VIP (Variable Importance of Projection) score >1 and regression coefficients to compute HCA. orthogonal partial least square regression discriminant analysis (OPLS-DA) showing differences among metabolic features under different levels of co-cultivation in (E) *Lb* & in (F) *Th.*

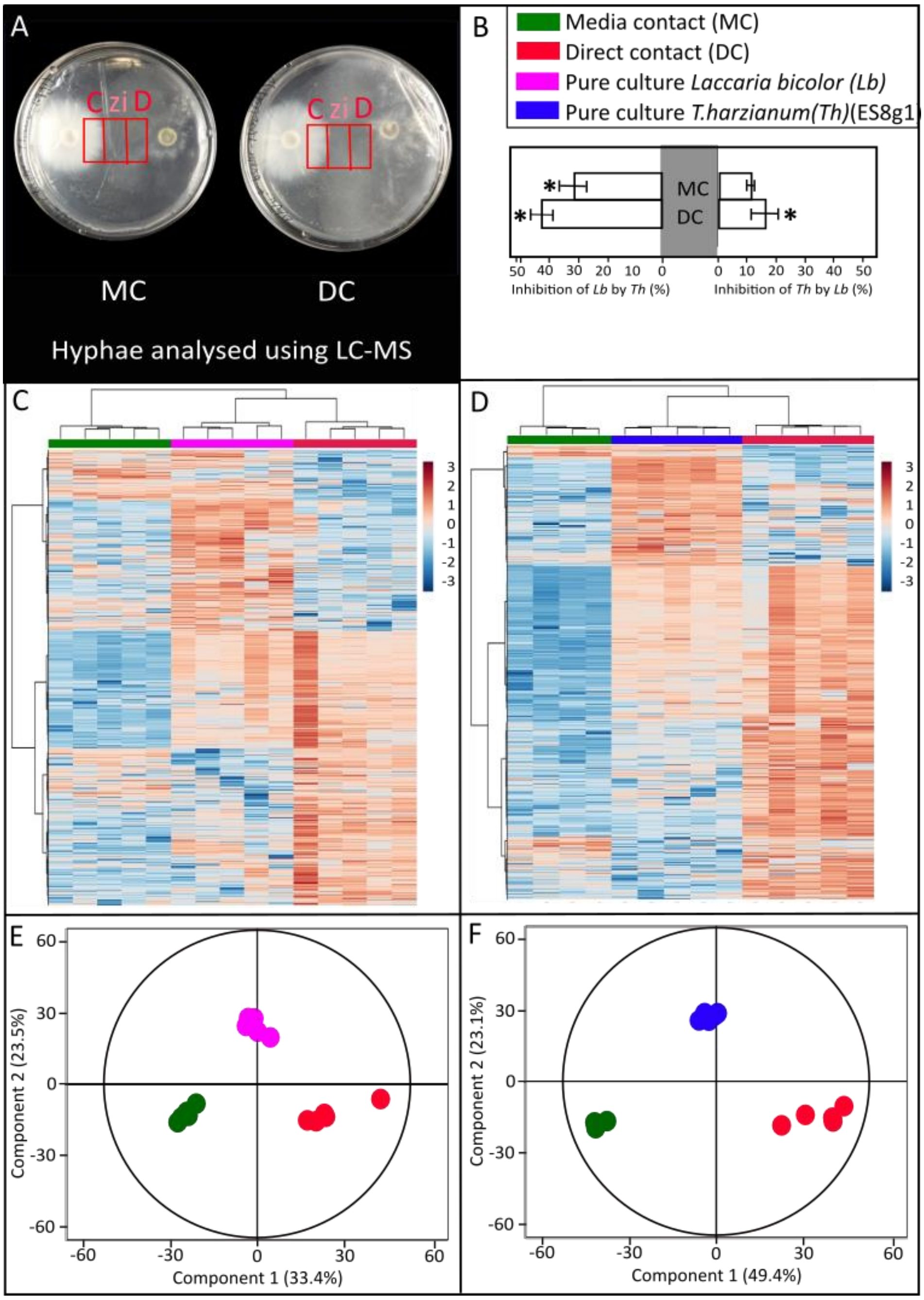

**Fig S6.** Metabolomic analysis of the hyphae from the co-cultivation experiment of *Laccaria bicolor* (*Lb*) and *T. harzianum (Th) (*ES8g1*).* (A) Exemplary image of the confrontation assay showing the three different zones of sampling of hyphae and media across media contact (MC) and direct contact (DC). (B) Growth inhibition of *Th* on *Lb* (left) and vice versa (right) under different levels of co-cultivation compared to pure cultures. Significances are denoted as asterisks (one-way ANOVA and Tukey HSD, $p < 0.05$); mean ± SE; Values are average of 5 replicates. Hierarchical clustering analysis of the peak area of features from cultures of (C) *Laccaria* and (D) *Th* hyphae grown in media contact (MC) direct contact (DC) and pure cultures. Features are selected based on having a VIP (Variable Importance of Projection) score >1 and regression coefficients to compute HCA. orthogonal partial least square regression discriminant analysis (OPLS-DA) showing differences among metabolic features under different levels of co-cultivation in (E) *Lb* & in (F) *Th*.

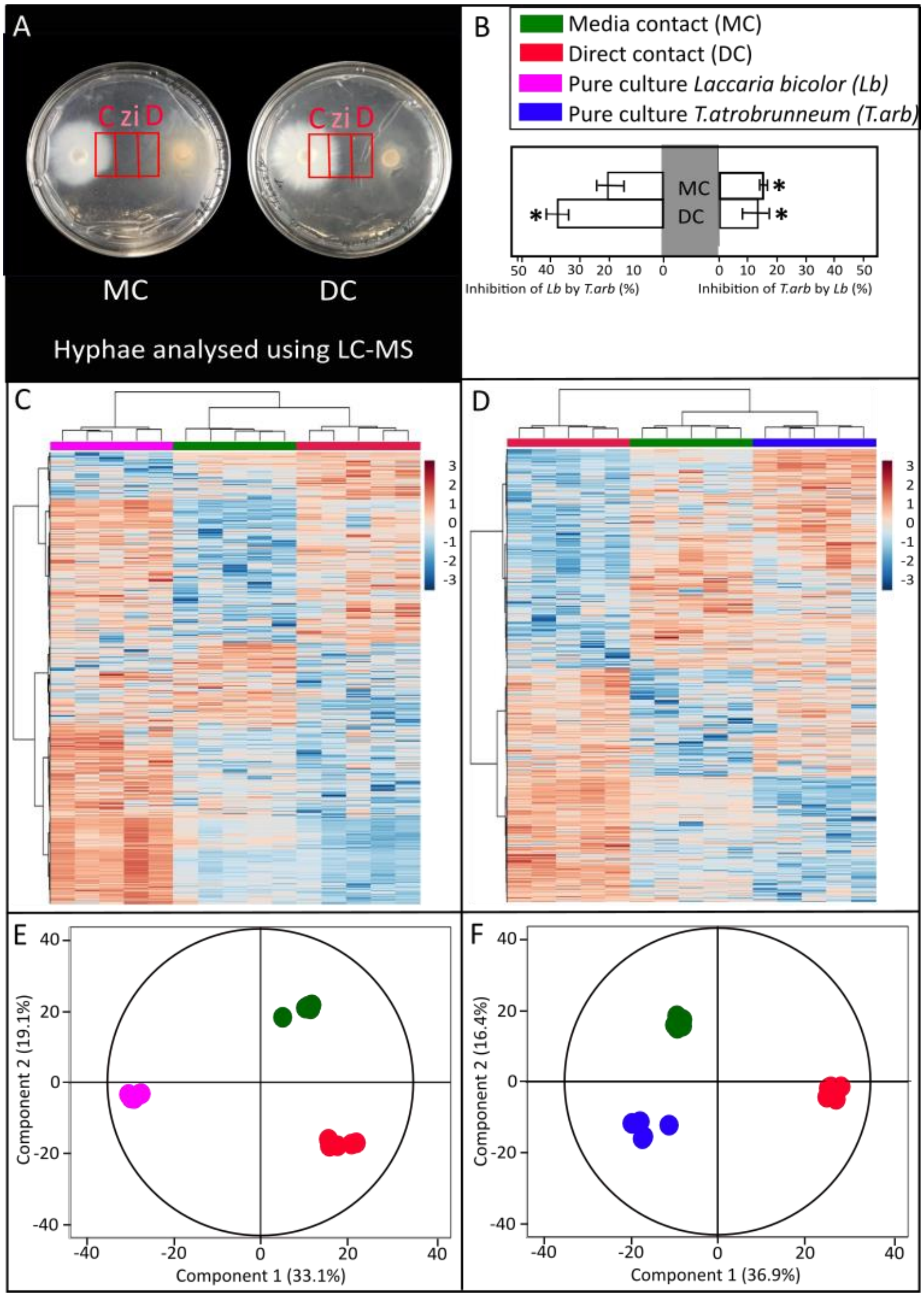

**Fig S7.** Metabolomic analysis of the hyphae from the co-cultivation experiment of *Laccaria bicolor* (*Lb*) and *T. atrobrunneum (T.arb).* (A) Exemplary image of the confrontation assay showing the three different zones of sampling of hyphae and media across media contact (MC) and direct contact (DC). (B) Growth inhibition of *T. arb* on *Lb* (left) and vice versa (right) under different levels of co-cultivation compared to pure cultures. Significances are denoted as asterisks (one-way ANOVA and Tukey HSD, $p < 0.05$); mean ± SE; Values are average of 5 replicates. Hierarchical clustering analysis of the peak area of features from cultures of (C) *Laccaria* and (D) *T. arb* hyphae grown in media contact (MC) direct contact (DC) and pure cultures. Features are selected based on having a VIP (Variable Importance of Projection) score >1 and regression coefficients to compute HCA. orthogonal partial least square regression discriminant analysis (OPLS-DA) showing differences among metabolic features under different levels of co-cultivation in (E) *Lb* & in (F) *T. arb.*

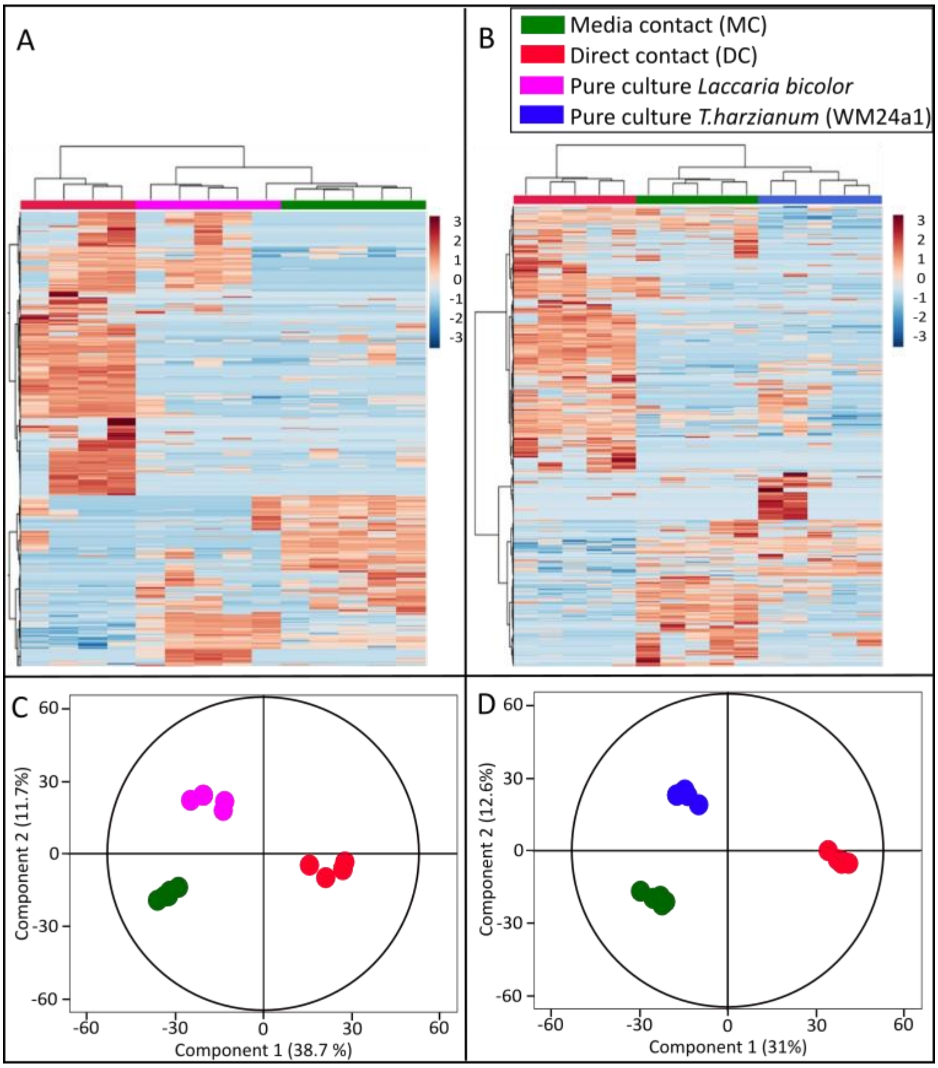

**Fig S8.** Metabolomic analysis of the media from the co-cultivation experiment of *Laccaria bicolor* (*Lb*) and *T. harzianum (*WM24a1*) (Th).* (A) Schematic representation of the co-cultivation experiment between *Lb* and *Th* showing the position of fungal inoculants from the two species, cellophane sheet, and the underlying media. (B) Experimental set-up of the confrontation assay showing the three different zones of sampling of media across media contact (MC) and direct contact (DC). Hierarchical clustering analysis of the peak area of features from cultures of (C) *Lb* and (D) *Th* grown also as media contact (MC) direct contact (DC) and pure cultures. Orthogonal partial least square regression discriminant analysis (OPLS-DA) showing differences among metabolic features in media under different levels of co-cultivation in (E) *Lb* & (F)*Th*.

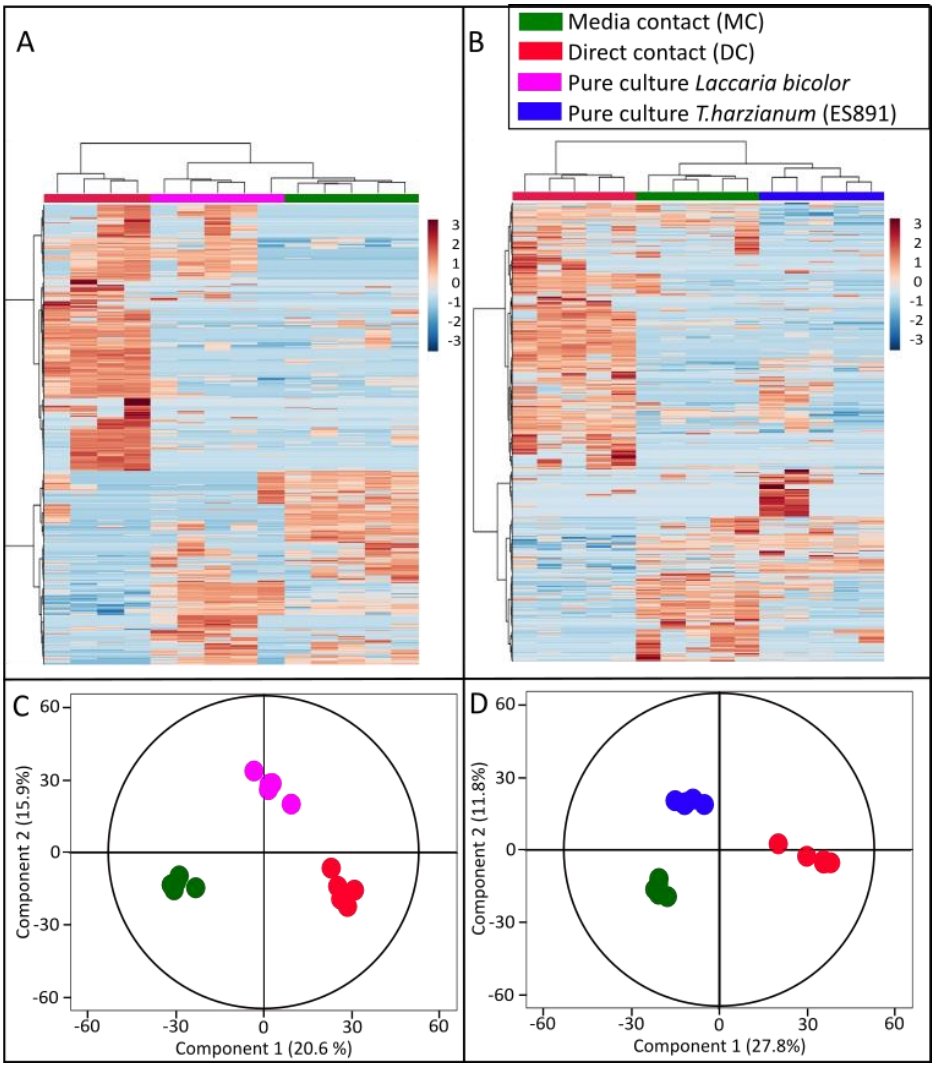

**Fig S9.** Metabolomic analysis of the media from the co-cultivation experiment of *Laccaria bicolor* (*Lb*) and *T. harzianum (*ES8g1*) (Th).* (A) Schematic representation of the co-cultivation experiment between *Lb* and *Th* showing the position of fungal inoculants from the two species, cellophane sheet, and the underlying media. (B) Experimental set-up of the confrontation assay showing the three different zones of sampling of media across media contact (MC) and direct contact (DC). Hierarchical clustering analysis of the peak area of features from cultures of (C) *Lb* and (D) *Th* grown also as media contact (MC) direct contact (DC) and pure cultures. Orthogonal partial least square regression discriminant analysis (OPLS-DA) showing differences among metabolic features in media under different levels of co-cultivation in (E) *Lb* & (F)*Th*.

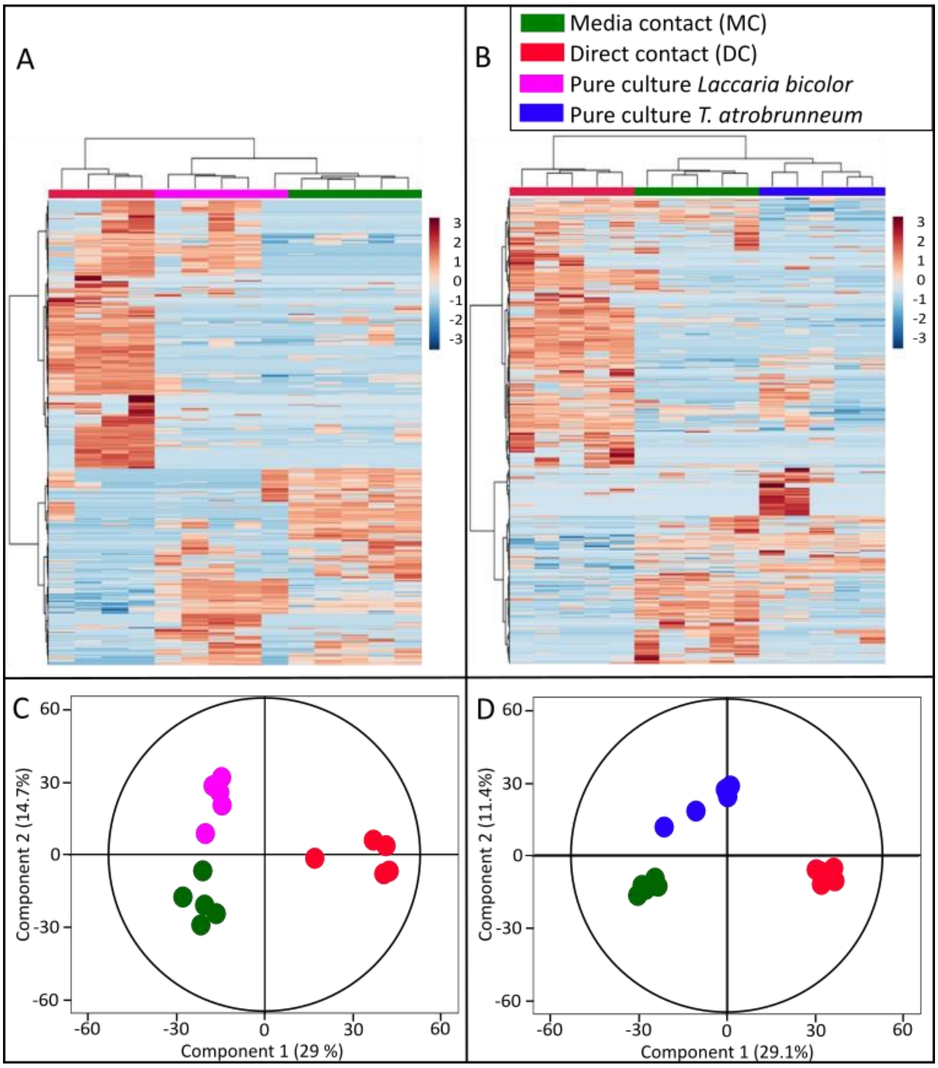

**Fig S10.** Metabolomic analysis of the media from the co-cultivation experiment of *Laccaria bicolor* (*Lb*) and *T. atrobrunneum (T.arb).* (A) Schematic representation of the co-cultivation experiment between *Lb* and *Th* showing the position of fungal inoculants from the two species, cellophane sheet, and the underlying media. (B) Experimental set-up of the confrontation assay showing the three different zones of sampling of media across media contact (MC) and direct contact (DC). Hierarchical clustering analysis of the peak area of features from cultures of (C) *Lb* and (D) *Tarb* grown also as media contact (MC) direct contact (DC) and pure cultures. Orthogonal partial least square regression discriminant analysis (OPLS-DA) showing differences among metabolic features in media under different levels of co-cultivation in (E) *Lb* & (F)*Tarb*.

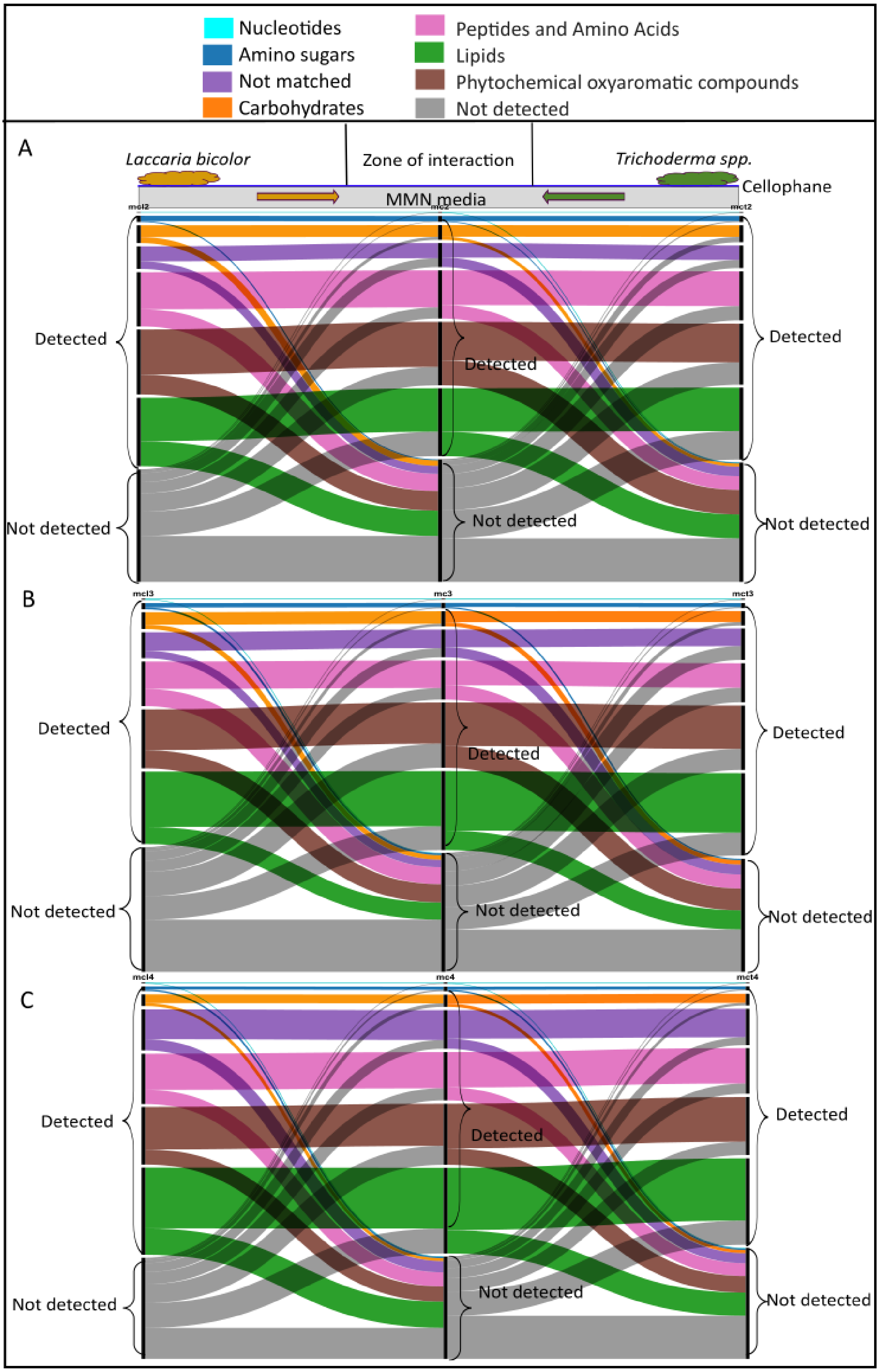

**Fig S11.** Alluvial plot showing the flow of exudates from *Laccaria bicolor* to *T. harzianum (*WM24a1*)* (A), *T. harzianum (*ES8g1*)* (B), *T. atrobrunneum* (C) and vice versa through the common growth media in media contact of the co-cultivation.

**Table S1**. Total emission strength of the different terpene classes across *Trichoderma* strains:

| CLASS | WM24a1 | WM24a1_AC | WM24a1_MC |
|---|---|---|---|
| **Monoterpenes** | 0 | 0 | 0 |
| **Oxygenated monoterpenes** | 0.272389 | 5.068172 | 2.179408 |
| **Sesquiterpenes** | 104.2598 | 109.8574 | 81.68247 |
| **Oxygenated sesquiterpenes** | 3.878956 | 4.437669 | 2.854758 |
| | **ES8g1** | **ES8g1_AC** | **ES8g1_MC** |
| **Monoterpenes** | 0.073041 | 3.118876 | 2.621989 |
| **Oxygenated monoterpenes** | 0.190644 | 0.298805 | 0.151202 |
| **Sesquiterpenes** | 95.04198 | 52.58572 | 76.39873 |
| **Oxygenated sesquiterpenes** | 3.215387 | 1.537527 | 2.291705 |
| | **MS8a1** | **MS8a1_AC** | **MS8a1_MC** |
| **Monoterpenes** | 14.14735 | 7.112237 | 9.910094 |
| **Oxygenated monoterpenes** | 0 | 0 | 0 |
| **Sesquiterpenes** | 23.37458 | 6.258328 | 10.02625 |
| **Oxygenated sesquiterpenes** | 0.625727 | 0.437164 | 0.630441 |
| | ***T.arb*** | ***T.arb_AC*** | ***T.arb_MC*** |
| **Monoterpenes** | 21.73812 | 34.28836 | 7.456772 |
| **Oxygenated monoterpenes** | 0.481796 | 0.632981 | 0.513673 |
| **Sesquiterpenes** | 131.9829 | 56.06847 | 49.19101 |
| **Oxygenated sesquiterpenes** | 3.624867 | 1.5577 | 1.348774 |
| | ***Laccaria bicolor*** | | |

| | |
|---|---|
| **Monoterpenes** | 13.40648 |
| **Oxygenated monoterpenes** | 0 |
| **Sesquiterpenes** | 0.483584 |
| **Oxygenated sesquiterpenes** | 0 |

**Table S2**. List of internal standards used for normalisation:

| Internal standard | Neutral Mass | Mode |
|---|---|---|
| **Bergapten** | 216.04 | + |
| **Plumbagin** | 188.05 | + |
| **Dihydrocaffeic acid** | 182.06 | - |
| **3,4-dihydromandelic acid** | 184.04 | - |

**Table S3**. LC-MS data processing parameters in Metaboscape 4.0:

| **Peak detection** | | |
|---|---|---|
| **Intensity threshold/spectra** | | 1000 |
| **Minimum peak length /spectra** | | 7 |
| **Feature signal** | | Area |
| **Recursive Feature extraction** | | |
| **Minimum peak length recursive/spectra** | | 6 |
| **Minimum number of features for extraction** | | 2/5 |
| **Presence of features in minimum number of analyses** | | 2/5 |
| **MS/MS import method** | | Average |
| **EIC correlation** | | 0.7 |
| **Ionization mode** | **Positive mode** | **Negative mode** |
| **Primary ion** | $[M+H]^+$ | $[M-H]^-$ |
| **Seed ions** | $[M+Na]^+$,$[M+K]^+$ | $[M+Cl]^-$ |
| **Common ions** | $[M-H20+H]^+$, | $[M-H20+H]^-$, |

| | | |
|---|---|---|
| | $[2M+H]^+$,<br>$[M+H+CH3CN]^+$ | $[2M+H]^-$,<br>$[M-H+CH3CN]^-$<br>$[M-H+HCOOH]^-$ |

**Table S4**. Differentially regulated metabolic features in hyphae of the fungal strains under different cocultivation scenarios:

| ***Trichoderma* strains on contact with *Laccaria*** | | | | | | | | |
|---|---|---|---|---|---|---|---|---|
| | **WM24a1** | | **ES8g1** | | **MS8a1** | | ***T. atrobrunneum*** | |
| | **Up** | **Down** | **Up** | **Down** | **Up** | **Down** | **Up** | **Down** |
| **MC** | 60 | 314 | 114 | 511 | 5 | 488 | 135 | 79 |
| **DC** | 162 | 300 | 595 | 522 | 159 | 319 | 434 | 332 |
| ***Laccaria* on contact with *Trichoderma* strains** | | | | | | | | |
| **MC** | 155 | 444 | 71 | 409 | 22 | 401 | 430 | 103 |
| **DC** | 126 | 339 | 102 | 235 | 196 | 254 | 110 | 395 |

**Table S5**. Differentially regulated metabolic features in the exudates of the fungal strains under different cocultivation scenarios:

| ***Trichoderma* strains on contact with *Laccaria*** | | | | | | | | |
|---|---|---|---|---|---|---|---|---|
| | **WM24a1** | | **ES8g1** | | **MS8a1** | | ***T. atrobrunneum*** | |
| | **Up** | **Down** | **Up** | **Down** | **Up** | **Down** | **Up** | **Down** |
| **MC** | 25 | 14 | 3 | 60 | 70 | 22 | 3 | 11 |
| **DC** | 16 | 26 | 11 | 19 | 112 | 47 | 4 | 6 |
| ***Laccaria* on contact with *Trichoderma* strains** | | | | | | | | |
| **MC** | 31 | 74 | 49 | 29 | | 15 | 9 | 50 |
| **DC** | 16 | 77 | 12 | 89 | 19 | 59 | 20 | 76 |
